# Export of diverse and bioactive peptides through a type I secretion system

**DOI:** 10.1101/2023.01.26.525739

**Authors:** Sun-Young Kim, Jennifer K. Parker, Monica Gonzalez-Magaldi, Mady S. Telford, Daniel J. Leahy, Bryan W. Davies

**Affiliations:** Department of Molecular Biosciences, The University of Texas at Austin, Austin, Texas; John Ring LaMontagne Center for Infectious Diseases, The University of Texas at Austin, Austin, Texas

**Keywords:** Secretion, Peptide, Gram-negative bacteria, T1SS

## Abstract

Microcins are peptide antibiotics secreted by Gram-negative bacteria that inhibit the growth of neighboring microbes. They are exported from the cytosol to the environment in a one-step process through a specific type I secretion system (T1SS). While the rules governing export of natural or non-native substrates have been resolved for T1SSs that secrete large proteins, relatively little is known about substrate requirements for peptides exported through T1SSs that secrete microcins. Here, we investigate the prototypic microcin V T1SS from *Escherichia coli* and show it can export a remarkably wide range of natural and synthetic peptides. We demonstrate that secretion through this system is not affected by peptide charge or hydrophobicity and appears only constrained by peptide length. A varied range of bioactive peptides, including an antibacterial peptide, a microbial signaling factor, a protease inhibitor, and a human hormone, can all be secreted and elicit their intended biological effect. Secretion through this system is not limited to *E. coli*, and we demonstrate its function in additional Gram-negative species that can inhabit the gastrointestinal tract. Our findings uncover the highly promiscuous nature of peptide export thorough the microcin V T1SS, which has implications for native cargo capacity and use of Gram-negative bacteria for peptide research and delivery.

**Importance:** Microcin type I secretion systems in Gram-negative bacteria transport antibacterial peptides from the cytoplasm to the extracellular environment in single step. In nature, each microcin secretion system is generally paired with a specific peptide. We know little about the export capacity of these transporters and how peptide sequence influences secretion. Here, we investigate the microcin V type I secretion system. Remarkably, our studies show this system can export diverse peptides and is only limited by peptide length. Furthermore, we demonstrate that various bioactive peptides can be secreted, and this system can be used in Gram-negative species that colonize the gastrointestinal tract. These finding expand our understanding of secretion through type I systems and their potential uses in peptide applications.

## Introduction

Gram-negative type I secretion systems (T1SSs) are unique in their ability to translocate proteinaceous cargos from the cytoplasm to the extracellular environment in one step (1, 2). While commonly described as one group, T1SSs can be subdivided into those that secrete large protein cargos and those that secrete small peptide cargos (3). These two subgroups differ in their method of processing of cognate substrates and suggested models for secretion (3, 4).

T1SSs are best known for export of large protein toxins and enzymes (2, 4). A well-studied example is the *Escherichia coli* hemolysin A T1SS, which secretes the hemolytic toxin, hemolysin A (HlyA) (2–4). The protein cargos encode a C-terminal signal peptide (SP) that is recognized by their cognate T1SSs. Following SP recognition, proteins are extruded linearly through a membrane tunnel to the environment, where they fold for action (5). The SP is retained by the exported protein. Overall charge, isoelectric point, and folding rate of the cargo can each influence protein secretion efficiency (1, 6, 7). This knowledge helps us understand sequence constraints on natural protein cargos and provides insight on how these systems may be co-opted for production and delivery of heterologous proteins (1, 8–11).

The lesser studied subgroup of Gram-negative T1SSs specializes in the export of small (< 10 kDa) peptide antibiotics, called microcins (12, 13). These T1SSs are distinct in their secretion mechanism, recognizing and cleaving a short, N-terminal SP during cargo export. The most well-studied example is the microcin V (MccV, previously referred to as colicin V) T1SS. The MccV system was identified in *E. coli* (14–17) and is comprised of three parts: a C39 peptidase-containing ATP-binding cassette transporter (PCAT), CvaB, a membrane fusion protein (MFP), CvaA, and an outer membrane efflux protein (OMP), TolC (17) (Fig. 1A). MccV is synthesized as a 103-amino-acid precursor peptide containing an N-terminal 15-amino-acid SP. The peptidase domain (PEP) of CvaB precisely cleaves the SP in an ATP-dependent manner, releasing the mature MccV for transit to the extracellular environment (18). One model for PCAT-driven peptide export is an alternating-access transport pathway (19, 20), which is not a generally believed model for linear extrusion of large protein cargo such as HlyA (3, 4).

**Figure 1.**
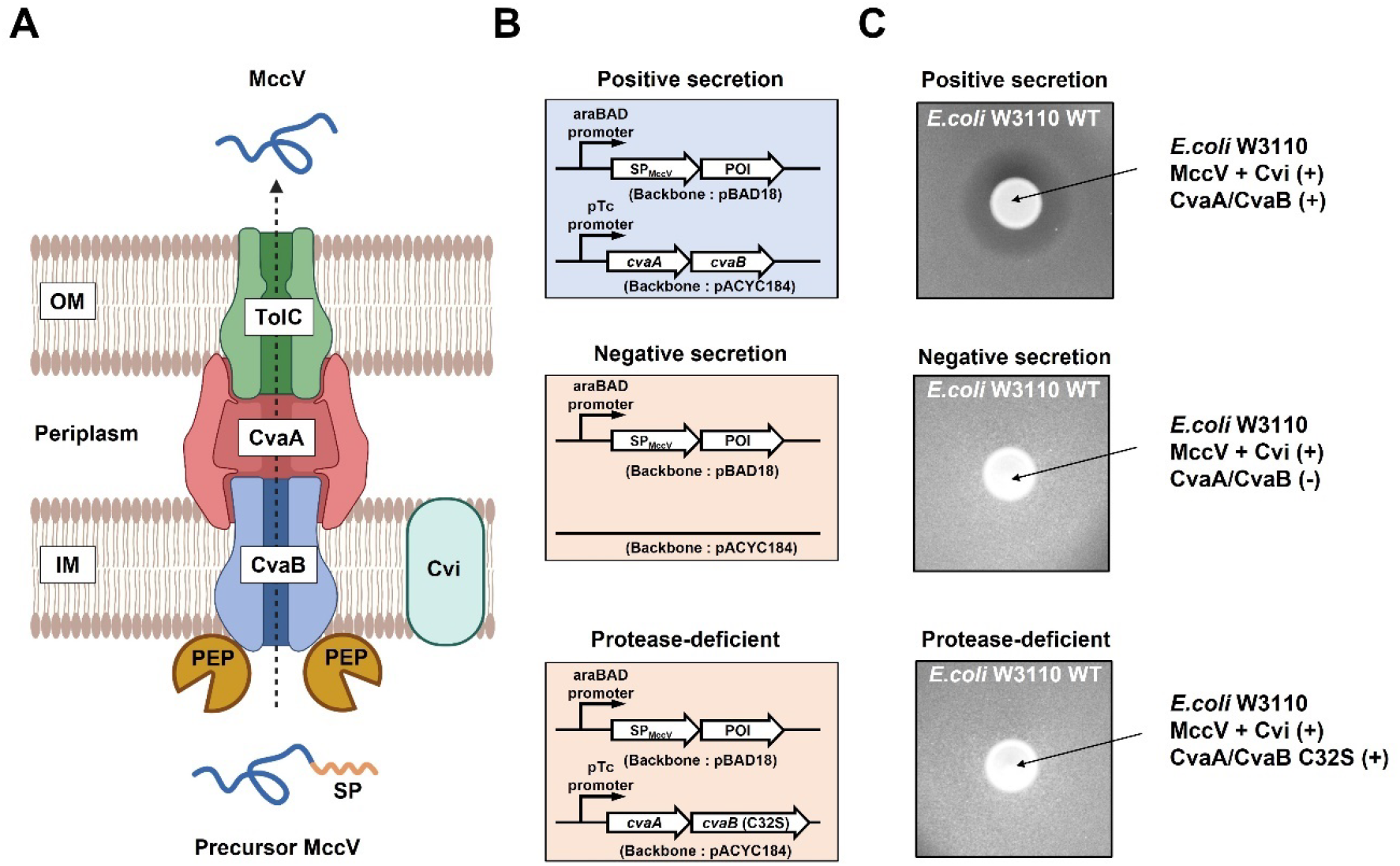
Construction of the secretion system. (A) The MccV secretion complex is composed of CvaA, CvaB, and TolC, and forms a channel for movement of cargo peptides from the cytoplasm to the extracellular environment. The 15-amino-acid signal peptide (SP) sequence of MccV (SP_MccV_) is cleaved by the peptidase domain (PEP) of CvaB during export. Cvi is an immunity protein that protects host bacterial cells from secreted MccV. This figure was created with BioRender.com. (B) Two-plasmid peptide secretion system is shown. Positive secretion strains encode a plasmid (pBAD18) expressing SP_MccV_ conjugated to the peptide of interest (POI) and a plasmid (pACYC184) expressing CvaA and CvaB. Control strains encode the same POI but carry either empty pACYC184 with no CvaAB (negative secretion) or pACYC184 expressing CvaA/CvaB C32S (protease-deficient secretion). Our secretion system uses chromosomally expressed TolC or homologs. (C) The results of MccV zone of inhibition assays are shown. *E. coli* W3110 expressing MccV and Cvi were spotted on a lawn of susceptible *E. coli* W3110. Only the positive secretor generated a zone of inhibition. All samples were spotted on the sample agar plate. The result is representative of biological triplicate.

Despite their importance in bacterial competition (21), only 10 microcins and their cognate T1SSs have been identified among Gram-negative bacteria (12, 13). The mature peptides of these 10 microcins share sequence similarities including high glycine content (>12%) and high hydrophobic residue content (>50%). Previous reports indicate that the MccV system can secrete MccV and related peptide antibiotics (22–24), suggesting export is not limited to the cognate cargo. However, it is unknown how the physical and chemical features of peptides might influence their export through microcin T1SSs. This lack of information limits our understanding of constraints on microcin sequence evolution and the potential cargos capable of export through microcin T1SSs.

Here, we show that the MccV system is a versatile peptide secretion system capable of exporting a diverse range of synthetic and bioactive peptides. Remarkably, secretion is not dependent on peptide physicochemical properties and appears only limited by peptide length. Furthermore, the MccV system is functional in several related species of Gram-negative bacteria adapted to gut colonization. Our work provides the first comprehensive study of cargo requirements for export through the MccV system and suggests its potential as a general peptide secretory platform in Gram-negative bacteria.

## Results

### Development of a two-plasmid peptide secretion system

A model of the native microcin V (MccV) type I secretion system (T1SS) is illustrated in Fig. 1A. CvaB encodes a C39 peptidase domain (PEP) that cleaves the 15-amino-acid signal peptide (SP) of pre-MccV (16, 25, 26). MccV then proceeds through CvaA and TolC as it exits the cell to reach the extracellular space. An immunity protein (Cvi) is co-expressed with MccV and protects the host cell against MccV’s antibacterial action (15). In nature, CvaAB, MccV, and Cvi are encoded on the same plasmid, and TolC is expressed from the chromosome (14, 15).

To investigate peptide features that influence export through a microcin T1SS, we developed an MccV secretion system that separates export functions from export cargo (Fig. 1B). The export components, CvaAB, are encoded on pACYC184 and are constitutively expressed. Peptides designed for secretion are cloned as fusion sequences with the MccV signal peptide (SP_MccV_) on pBAD18 under arabinose-inducible control. We refer to positive secretion (PS) if the bacteria carry plasmids encoding the SP_MccV_-conjugated peptide of interest (POI) and the wild-type (WT) CvaAB export system. Negative secretion (NS) refers to control bacteria that contain the same plasmids for expression of the SP_MccV_-POI, but with pACYC184 lacking CvaAB (empty vector). It was previously shown that a C32S mutation in CvaB abolished secretion of MccV due to disruption of the protease function of CvaB (18, 27). Protease-deficient secretion (PD) refers to bacteria that contain pBAD18 for expression of the SP_MccV_-POI and pACYC184 expressing CvaA with a protease-deficient CvaB mutant (C32S). Both negative and protease-deficient secretion strains act as controls to ensure that observed peptide secretion is due to CvaAB activity and not due to bacterial cell lysis in our experiments.

To ensure our two-plasmid approach enables robust cargo secretion, we expressed native MccV along with its immunity protein (Cvi) in *E. coli* W3110. When plated on a lawn of sensitive *E. coli*, a zone of inhibition (ZOI) can be observed around our microcin-secreting strain. We observed that only *E. coli* encoding MccV, Cvi, and WT CvaAB produced a visible ZOI against susceptible *E. coli* (Fig. 1C). Strains encoding MccV, Cvi, and empty pACYC184 or the CvaB mutant did not form a ZOI, indicating they could not secrete MccV.

We next generated a C-terminal V5 epitope-tagged MccV (MccV_V5) to track its secretion from *E. coli* W3110 by immunoblot. Two glycine residues (GG linker) were included between MccV and the V5 tag to allow for flexibility of the epitope. This MccV_V5 fusion retained its inhibitory activity (Fig. 2A). Only *E. coli* encoding MccV_V5, Cvi, and WT CvaAB could inhibit susceptible *E. coli* (Fig. 2A). This result indicates that MccV_V5 is secreted and shows the same dependency on CvaAB as the native MccV.

**Figure 2.**
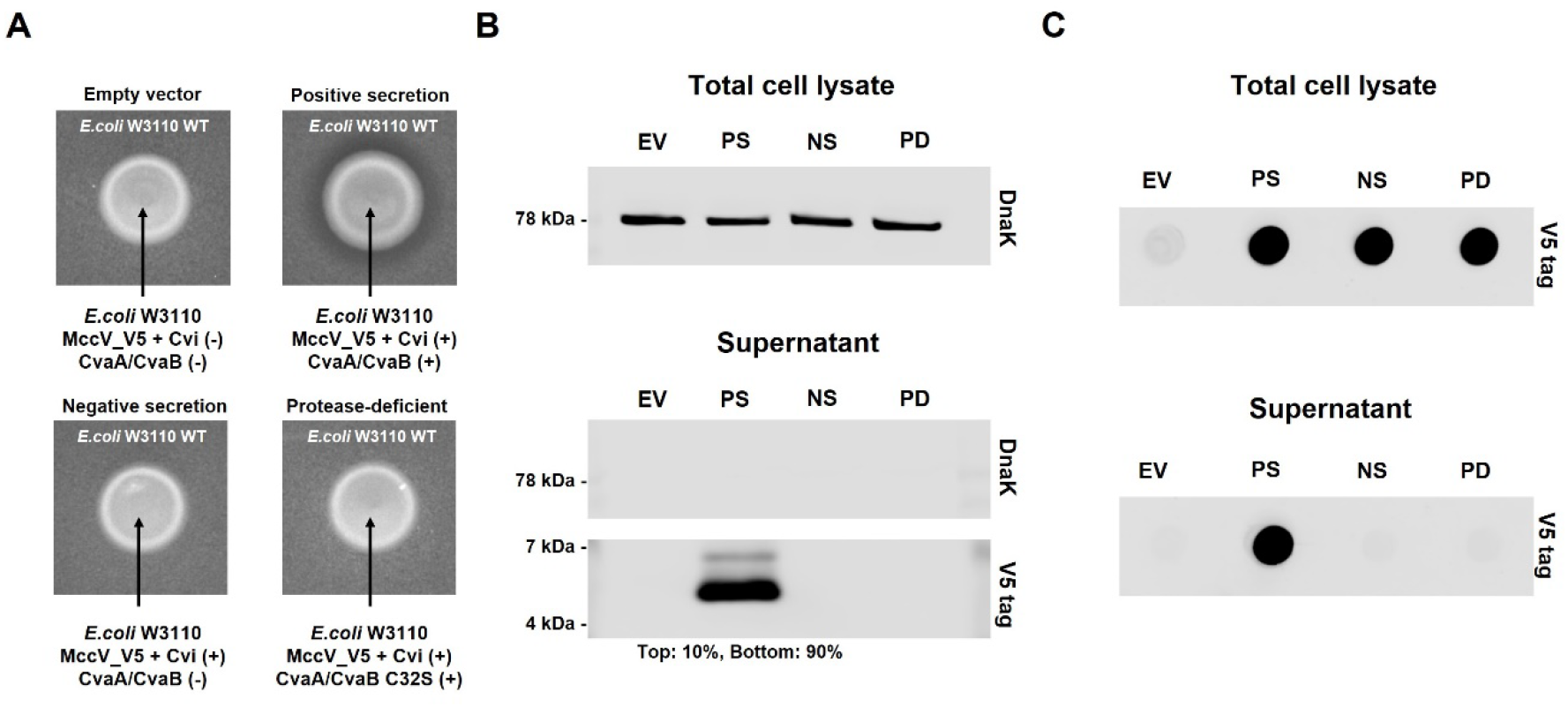
Recombinant peptide secretion via the MccV system. (A) Zone of inhibition assays were performed as described in Fig. 1C. Empty vector (EV) strain carries empty plasmids, pBAD18 and pACYC184. Positive secretion (PS) strain encodes MccV_V5, Cvi, and CvaAB. Negative secretion (NS) strain encodes MccV_V5, Cvi and empty pACYC184. Protease-deficient secretion (PD) strain encodes MccV_V5, Cvi and CvaA/CvaB C32S. All samples were spotted on the same agar plate. (B) Western blot detecting secreted MccV_V5 from *E. coli* W3110 is shown. Culture supernatant or pellet (total cell lysate) were directly suspended in sample buffer and loaded into each well. Antibody targets are described on the right side. The percentages of V5 tag top band and bottom band intensity from PS supernatant sample is shown at the bottom. (C) The result of dot blots against the V5 tag in total cell lysate and supernatant is shown. Samples from all strains (EV, PS, NS, and PD) were loaded into wells of the dot blot apparatus and detected with anti-V5 antibody. The results are representative of biological triplicates and all western or dot blot images were prepared from a single membrane.

We induced expression for two hours and investigated the supernatant for the presence of MccV_V5. In supernatant from our PS strain, we detected a dominant band migrating near 5–6 kDa (Fig. 2B), which is consistent with previous observations of MccV (16, 17, 25). We observed a weaker band migrating at a slightly higher molecular weight that likely represents unprocessed precursor MccV_V5 (17). This suggests a small amount of peptide (~10%) can escape without N-terminal processing under these conditions. MccV_V5 was only detected from bacteria that encoded WT CvaAB (Fig. 2B). This result is consistent with the ZOI results in Fig. 2A and further supports the dependence of peptide secretion on CvaAB. Cytoplasmic protein, DnaK, was not observed in any of the supernatant samples, but was readily observed in total cell lysate, indicating that bacteria were not lysing during secretion of MccV_V5. To simplify the process for detecting secreted peptides, we performed dot blot of MccV_V5 from supernatants and total cell lysates (Fig. 2C). While MccV_V5 was detected in all total cell lysates, it was only present in the supernatant of our PS strain, consistent with our western blot analysis (Fig. 2B). These results support the use of a two-plasmid platform and dot blotting to detect secreted peptides.

### Construction of a sequence-diverse synthetic peptide library

The MccV secretion system has only been shown to export MccV and related peptide antibiotics (22–24). It is unclear whether it can export other types of peptides and if there are biochemical or physical limitations on sequences that can be secreted. To begin to examine how chemistry and size influences peptide secretion through the MccV system, we generated a library of 40 random peptide sequences. Our library consists of four groups; each group includes ten peptides that have the same length (10, 20, 50 or 100 amino acids) but different, randomly generated sequences. Each peptide encodes a C-terminal V5 tag, which increases the size of Groups 1 - 4 to 26, 36, 66, and 116 amino acids respectively. Finally, each peptide encodes an N-terminal SP_MccV_ to direct export (Table S1). We hypothesized that using peptides of diverse composition and length would provide insight into the ability of the MccV system to secrete cargo peptides that vary from its original substrate, MccV (88 amino acids without the SP).

We plotted the distribution of charge and hydrophobicity for each peptide in our library (Fig. S1A) and the amino acid composition per group (Fig. S1B). For the latter, we analyzed amino acid sequences without the GG linker and V5 tag to avoid composition bias. Each of these measurements appeared well-distributed across the library as a whole. Group 1 encoded the shortest 26-amino-acid peptides and showed the narrowest distribution of charge and hydrophobicity. Based on this analysis, we deemed our library sufficiently random in length and chemistry to begin exploring peptide secretion.

### Secretion of a synthetic peptide library from *E. coli*

Each peptide in our library was expressed from *E. coli* W3110 with (PS) or without (NS) CvaAB to analyze their secretion levels. We performed dot blots to determine the relative amounts of each peptide in supernatants and total cell lysates after 8 hours of induction (Fig. 3A). To calculate secretion levels, each signal was first normalized by the cell density (OD_600_) of its respective culture to account for possible differences in bacterial abundance. Then, we subtracted the NS signal intensity to account for background signal. The differential signal intensity, or relative secretion level, of each peptide was graphed in Figure 3B and the distribution of secretion level is shown in Figure 3C.

**Figure 3.**
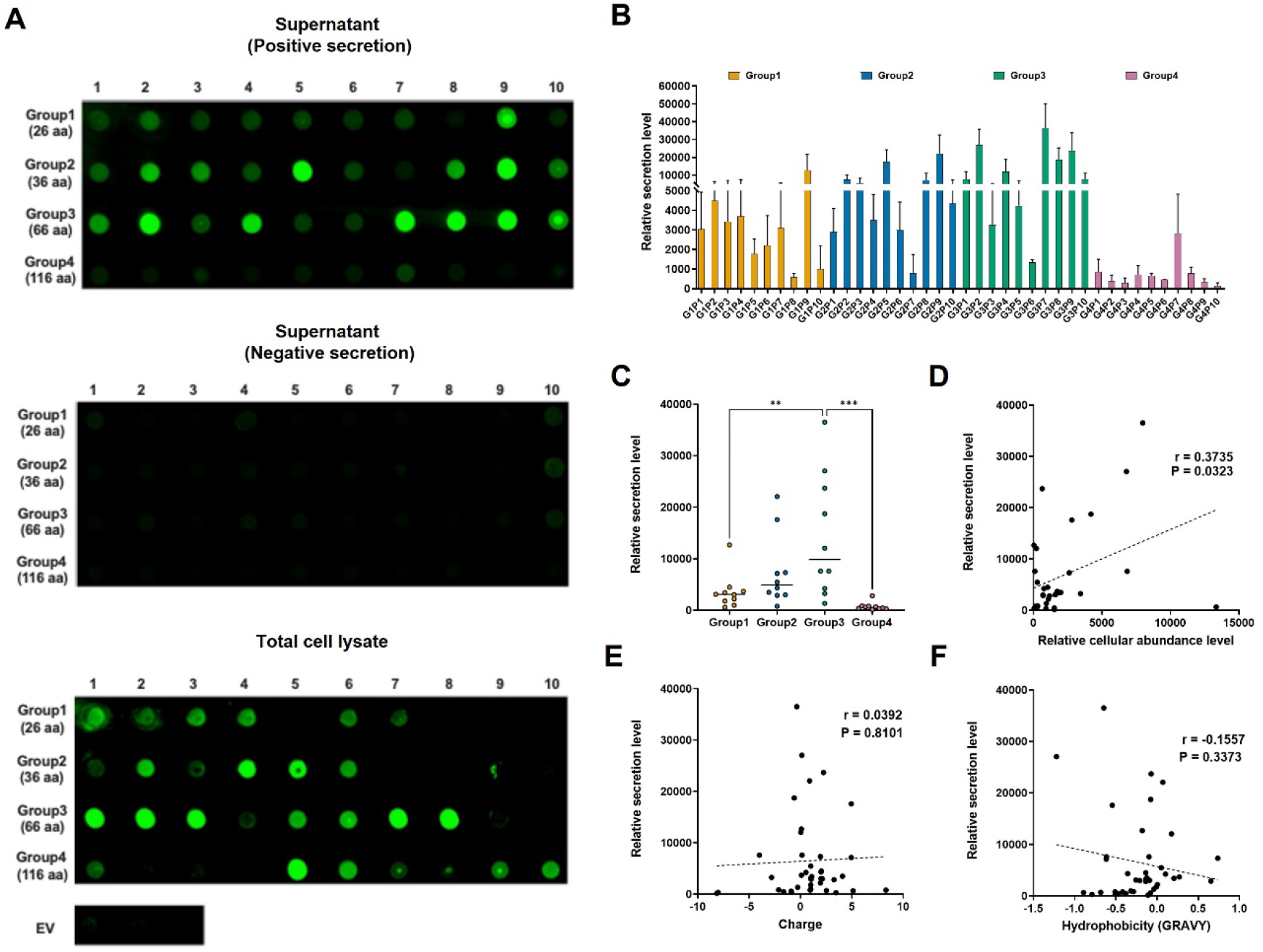
Secretion levels of random peptides. (A) The result of dot blots against the V5 tag from supernatant or total cell lysate samples of random peptides is shown. Each peptide group is shown on the left with the number of amino acids (aa), and the numbers on top represent each individual peptide in each group. Supernatant samples include both positive and negative secretion (no CvaAB). Total cell lysate samples were prepared by lysing negative secretion cultures. The results of total cell lysate samples were also shown with empty vector (EV) cell lysate samples. The results are representative of biological triplicates, and each image is from a single membrane. (B) Relative secretion levels are shown. Each group is represented as a different color; Group 1 (orange), Group 2 (blue), Group 3 (green), and Group 4 (purple). The mean of biological triplicate is shown with standard deviation. (C) The distribution of the mean secretion level of each group is shown. The median secretion level of each group is shown as a bar. Significant differences between groups are shown as asterisks (** = *P* < 0.01, *** = *P* < 0.001). Adjusted *P*-values were calculated by ANOVA with Tukey HSD test. (D, E, F) Correlation analysis between the mean of relative secretion level and (D) the mean of relative cellular abundance, (E) charge, and (F) hydrophobicity was performed. Grand average of hydropathicity index (GRAVY) is used to represent the hydrophobicity value of a peptide. Correlation coefficient (r value) and *P*-value of each analysis is shown. In the correlation assay for relative secretion level vs. relative cellular abundance level, we did not include the seven peptides (G1P5, G1P10, G2P7, G2P8, G2P9, G2P10, and G4P3) in which calculated cellular abundance levels were below zero.

Remarkably, and despite having little sequence relationship to the native cargo, all peptides in groups 1–3 (26–66 amino acids) were secreted well above background level and, in general, secretion levels increased from groups 1–3 (Fig. 3B, C). Interestingly, secretion levels abruptly dropped in group 4 (116 amino acids) as a whole (Fig. 3A, B). The near background level of secretion of this group of peptides suggests the MccV system has a maximum length preference somewhere between its native substrate size (MccV = 88 amino acids) and 116 amino acids. However, G4P7 (group 4 peptide 7) did show a modest level of secretion, indicating that length may not be a strict limitation.

In addition to length, we questioned if other peptide properties influenced their secretion. We first tested if global chemical characteristics of charge or hydrophobicity related to peptide secretion level. However, we found only negligible correlations between peptide charge (r = 0.0392, *P* > 0.05) and hydrophobicity (r = −0.1557, *P* > 0.05) with secretion levels across in our library (Fig. 3E and F), suggesting these features do not strongly influence peptide export. We hypothesized that peptide cellular abundance may influence the amount of peptide secreted. If more substrate is present in the cytoplasm, there may be more chance for it to be exported. To obtain cellular abundance levels, we calculated signal intensity of each peptide from total cell lysates by dot blot (Fig. 3A Total cell lysate). Samples were normalized for cell density and the signal intensity of control cell lysate that expressed neither a cargo peptide nor secretion machinery protein (empty vector). There was a small positive relationship (r = 0.3735, *P* < 0.05) between relative cellular abundance level and secretion level (Fig. 3D), suggesting that cellular abundance level can have a modest effect on the amount of peptide secreted. As has been found for other secretion systems, the abundance of peptide exported through the microcin V T1SS will likely be influenced by additional factors specific to each unique peptide cargo (1). Overall, our results imply that length, but not charge, hydrophobicity, or cellular abundance, is the only peptide metric that consistently and strongly correlates with secretion abundance.

### Peptide length influences secretion

Results from Figure 3 suggest that longer peptides tend to have a higher secretion level (groups 1– 3), before reaching a length limitation (group 4). Based on these observations, we hypothesized that a group 1 peptide (26 amino acids) will have a higher secretion level once its length is increased to group 2 length (36 amino acids), and a group 3 peptide (66 amino acids) will have a lower secretion level when its length is increased to group 4 length (116 amino acids). To test this concept, we selected two peptides G1P6 (group 1 peptide 6) and G3P2 (group 3 peptide 2). We generated new peptides, G1P6_2X (36 amino acids) and G3P2_2X (66 amino acids), whose sequences were lengthened by adding a direct repeat of their respective peptide sequence (maintaining a single V5 tag) and maintains similar biochemical properties to the parent peptides. We measured their relative secretion and cellular abundance levels by dot blot (Fig. 4A). G1P6_2X had a significantly higher secreted level compared to G1P6 (Fig. 4B). Interestingly, in the absence of CvaAB, the cellular abundance of G1P6 and G1P6_2X was similarly low (Fig. 4C), but in the presence of CvaAB the cellular abundance of G1P6_2X was significantly higher than G1P6 (Fig. 4D). This suggests that the increased secretion level of G1P6_2X may be partly related its increased cellular abundance when CvaAB is present. The opposite result was observed when comparing secretion of G3P2 and G3P2_2X. In this case, the longer G3P2_2X peptide showed significantly less secretion than the shorter parent sequence (Fig. 4A and B).

**Figure 4.**
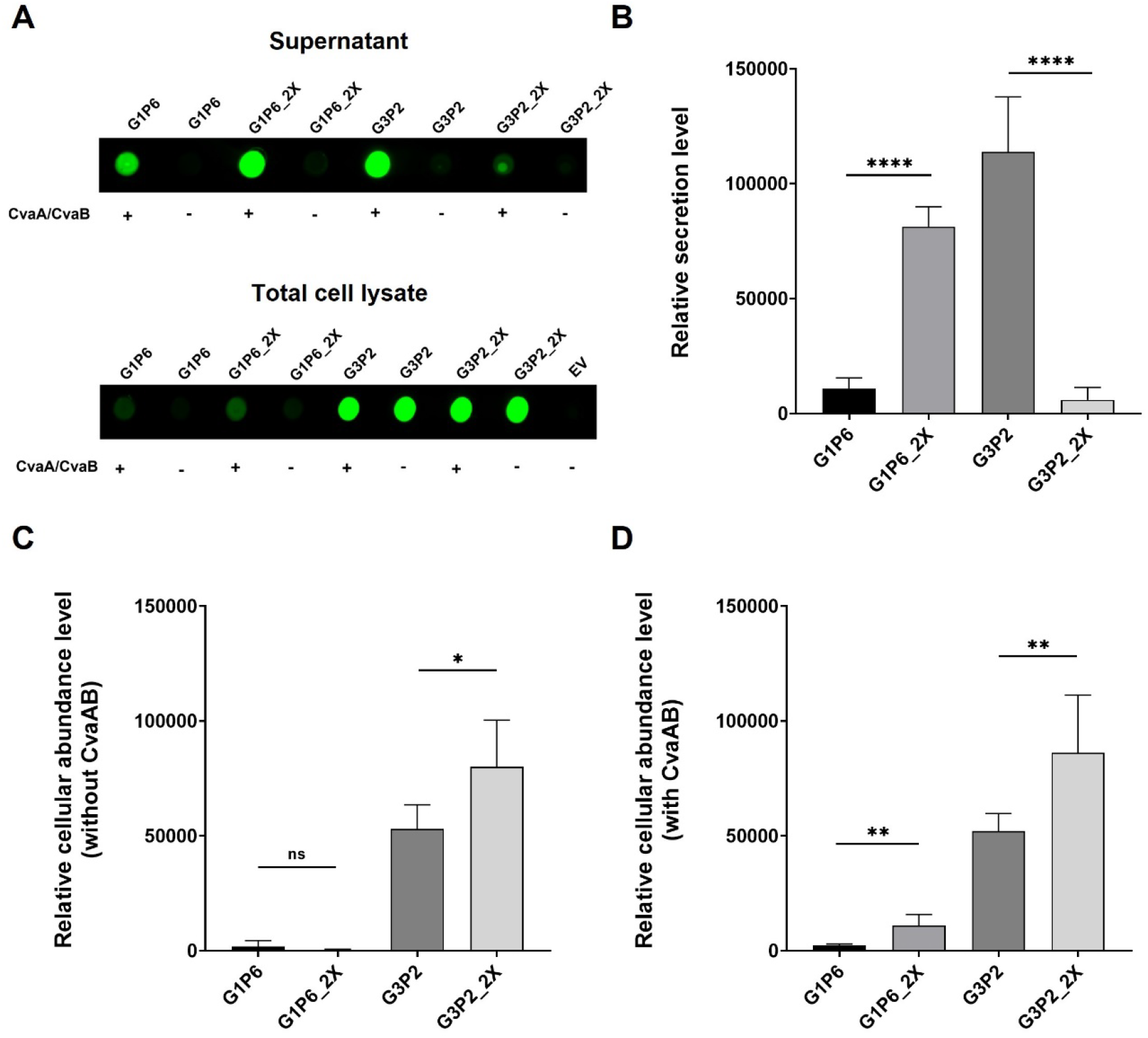
Size-dependent secretion via the MccV system. (A) Dot blot result showing V5 tag signal intensities of G1P6, G1P6_2X, G3P2, G3P2_2X supernatant and total cell lysate samples. V5 tag signal of empty vector (EV) total cell lysates is also shown. A representative dot blot image prepared from a single membrane is shown. (B, C and D) Relative secretion or cellular abundance levels of indicated peptides are graphed. The mean of six biological replicates is shown with standard deviation. Two-tailed *P*-values from unpaired t-test result is shown as ns (not significant, *P* > 0.05), or the number of asterisks (* = *P* < 0.05, ** = *P* < 0.01, **** = *P* < 0.0001).

However, G3P2_2X has higher cellular abundance regardless of the presence of the secretion apparatus (Fig. 4C and D). We hypothesized that G3P2_2X could not efficiently pass through the secretion machinery due to its longer length. Our results support the concept that the peptide cargo length is an important factor controlling the ability of a peptide to be secreted through the MccV secretion system.

### Heterologous sequences can be exported as efficiently as the native cargo

Our results suggest the MccV T1SS is promiscuous for cargo sequence and is only limited by peptide length. While a diverse set of sequences were exported (Fig. 3), it is unclear how well they were exported compared to the native cargo MccV. To investigate this question, we quantified the amount of V5-tagged natural substrate (MccV_V5) and a well-secreted peptide (Fig. 3B) from each of our library groups (G1P9, G2P9, G3P2, and G4P7). Supernatant from *E. coli* secreting MccV_V5, G1P9, G2P9, G3P2, and G4P7 was dot blotted after 8 hrs of induction (Fig. 5A). Secretion signal intensity was converted to mg/L per OD_600_ based on a standard curve generated using a commercially synthesized V5-tagged peptide (Fig. S2). G1P9, G2P9, and G3P2 supernatant concentrations were in the high μg/L to low mg/L (0.19–7.25 mg/L) per OD_600_ range (Fig. 5B). Interestingly, the concentrations of G3P2 and MccV_V5 were similar, suggesting that a heterologous peptide cargo can be secreted at a level comparable to the native cargo. As anticipated, G4P7 secretion level was low (Fig. 5A) and out of range of our standard curve. We also observed the same production trend as in Figure 3, where the order of abundance from highest to lowest was G3P2, G2P9, G1P9, and G4P7.

**Figure 5.**
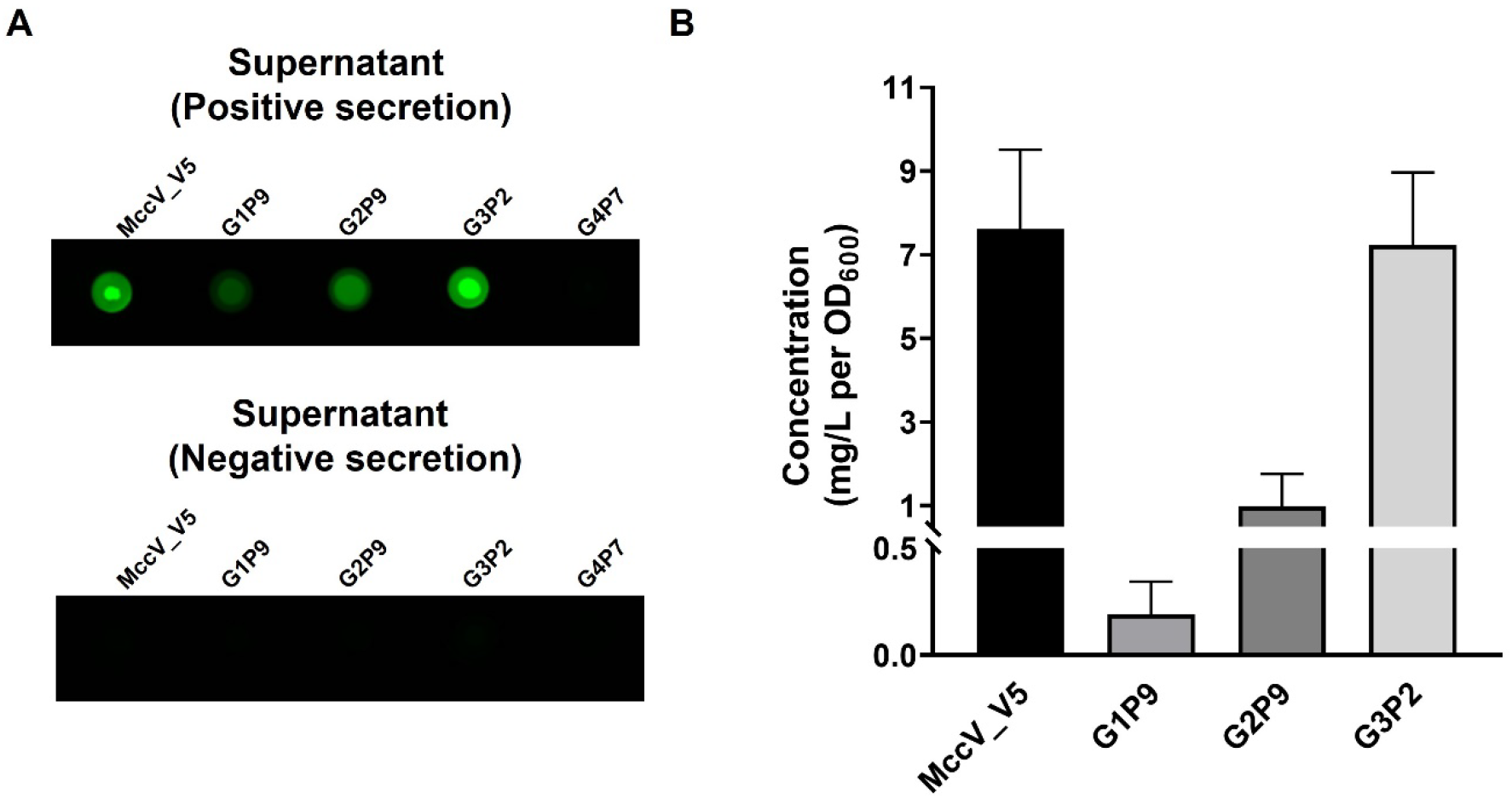
Secretion capacity of the MccV system. (A) Dot blot result showing V5 tag signal intensities of positive and negative secretion supernatants of selected peptides (MccV_V5, G1P9, G2P9, G3P2 and G4P7) is shown. A representative dot blot image prepared from a single membrane is shown. (B) Concentrations (mg/L per OD_600_) of selected peptides are shown as a graph. The mean of at least four biological replicates with standard deviation is shown.

### Abundance of secreted peptide is influenced by host strain and temperature

All our secretion tests so far had been conducted in *E. coli* K-12 strain W3110 at 37 °C. In Figure 3, we observed G1P5, G1P10, G2P6, G2P7, G3P3, and G3P5 have relatively lower secretion levels. We hypothesized that using a bacterial strain designed for protein production through deletion of cellular proteases (*E. coli* BL21(DE3)) and lowering growth temperature, which are general strategies for heterologous protein expression optimization (28), could increase peptide secretion levels. To test this hypothesis, peptide secretion levels from *E. coli* W3110 grown at 30 °C, *E. coli* BL21(DE3) grown at 37 °C, and *E. coli* BL21(DE3) grown at 30 °C were measured and normalized to levels produced under conditions used in Figure 3 (*E. coli* W3110 at 37 °C) (Fig. S3). Lowering the growth temperature and/or using BL21 cells significantly increased secretion levels of 5 out of the 6 peptides (G1P5, G2P6, G2P7, G3P3, and G3P5) from 2-17-fold. Only expression of G1P10 was not significantly increased. No single change (temperature or strain) consistently increased secretion. These results indicate that secretion of peptides through the MccV T1SS can be strongly influenced by environmental conditions and host strain, but the impact varies by cargo sequence.

### Secretion of bioactive peptides by the MccV system

Studies of non-native cargo export via the MccV system have focused on related peptide antibiotics.(22–24). To expand our knowledge, we sought to test the secretion of bioactive peptides from various organisms. We selected four peptides within the length constraints defined above that have diverse biological functions: pediocin PA-1, α-factor, eglin C, and epidermal growth factor (EGF). The functions and properties of the peptides are listed in Table S2. Each peptide was fused with the N-terminal SP_MccV_ to direct their export.

We began with pediocin PA-1, an antimicrobial peptide natively produced by the Gram-positive bacterium, *Pediococcus acidilactici*, and active against the Gram-positive bacterium *Listeria monocytogenes* (29). Similar to MccV (Fig. 1C), we performed ZOI assays by secreting pediocin PA-1 from *E. coli* W3110 spotted on a lawn of *L. monocytogenes*. When *E. coli* expressed both pediocin PA-1 and CvaAB, it inhibited *L. monocytogenes* growth and produced a ZOI (Fig. 6A). No ZOI was observed with our NS strain (no CvaAB) or the PD strain (CvaB C32S mutant), confirming the CvaAB-dependent secretion and action of pediocin PA-1. This indicates the MccV secretion system can produce and deliver heterologous peptide antibiotics.

**Figure 6.**
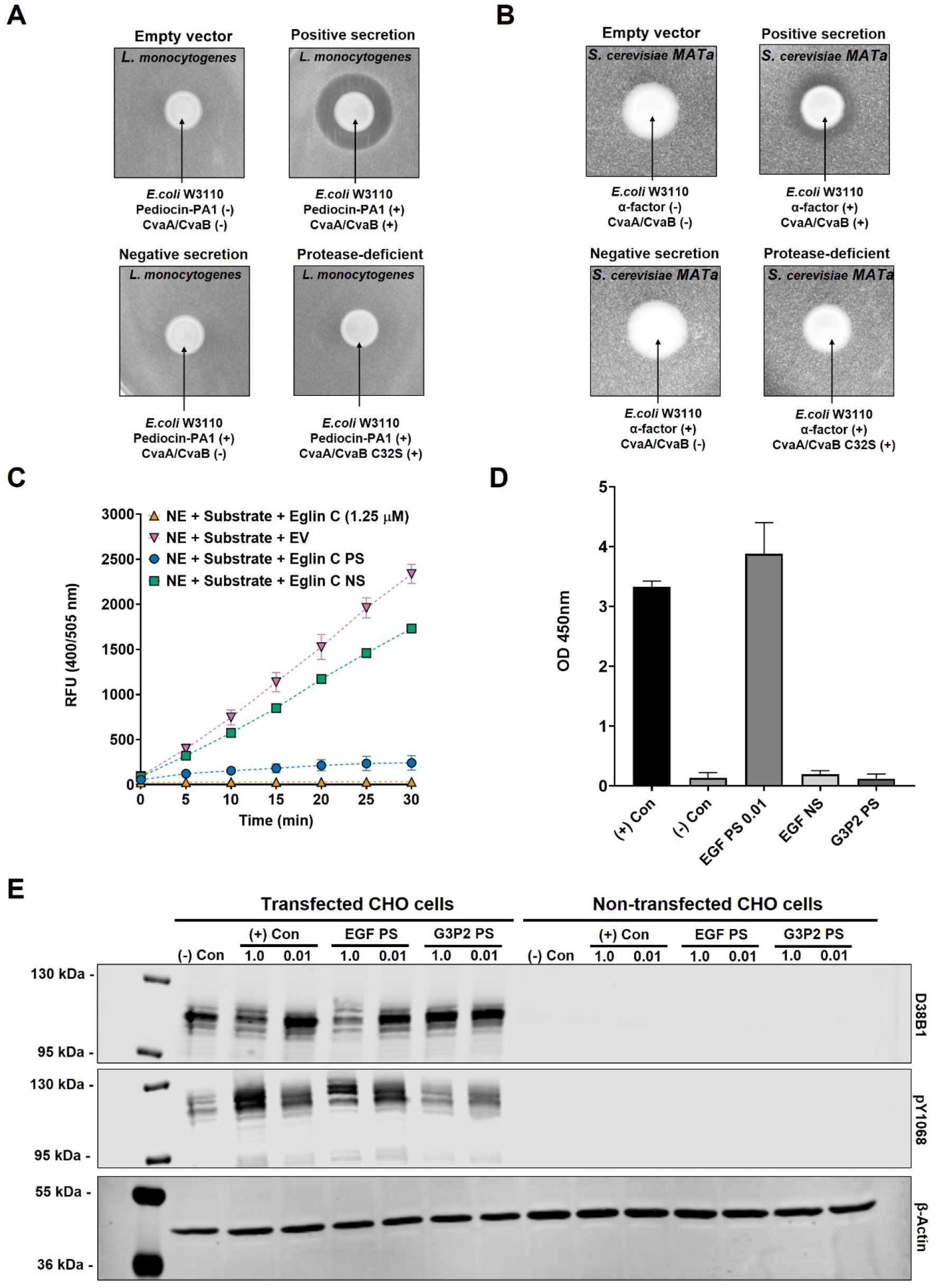
Bioactive peptides are secreted via the MccV system. (A and B) The results of two zone of inhibition assays are shown. The results are the representative images of biological triplicates. (C) The result of neutrophil elastase (NE) activity assay is shown. Relative fluorescence levels of samples at each time point are shown. All samples contain NE and substrate. 1.25 μM of recombinant eglin C, empty vector (EV) supernatant, eglin C positive secretion (PS) supernatant, or eglin C negative secretion (NS) supernatant was separately added into each sample. The mean of biological triplicate is plotted with standard deviation. (D) The result of ELISA for human epidermal growth factor (EGF) is shown. OD_450_ values indicate relative amounts of human EGF in samples. Positive control (+ Con) represents a sample containing standard EGF (1 ng/mL), Negative control (− Con) represents a sample containing assay buffer. EGF PS 0.01 represents 100-fold diluted EGF positive secretion supernatant in fresh media. EGF NS represents EGF negative secretion supernatant (no CvaAB). G3P2 PS represents G3P2 positive secretion supernatant. The mean of biological duplicate with standard deviation is shown. (E) The results of western blot assays are shown. Human EGFR (EGF receptor) transfected and non-transfected CHO cells were treated with each sample: (−) Con: untreated, (+) Con: standard EGF (100 ng/ml), EGF PS: EGF positive secretion supernatant, G3P2 PS: G3P2 positive secretion supernatant (non-specific peptide control). Cells were treated with either undiluted sample (1.0) or 100-fold diluted sample (0.01) in fresh media, and cell lysates were subjected to western blot against EGFR (D38B1), phosphorylated EGFR (pY1068) and β-Actin. The blots are representative images of biological duplicate.

We also used the ZOI test to assay α-factor production. α-factor is a pheromone released by *Saccharomyces cerevisiae* mating-type alpha cells that activates G-protein-coupled receptor (GPCR) Ste2p causing cell-cycle arrest in susceptible *S. cerevisiae* strains (MATa) (30). Figure 6B shows that, when *E. coli* W3110 expresses both α-factor and CvaAB, it inhibits the growth of susceptible *S. cerevisiae* MATa. This indicates the MccV system can secrete peptides active against GPCRs and impact evolutionarily distant organisms.

We next tested whether the MccV system secretes a metabolic inhibitor. Eglin C is a naturally occurring polypeptide from leeches and a potent inhibitor of serine proteases including elastase (31). Eglin C has been investigated for the treatment of gastrointestinal conditions (32). We tested the ability of secreted eglin C to inhibit neutrophil elastase (NE) compared to commercial eglin C. The assay measures cleavage of a substrate-fluorophore-conjugate by NE; hydrolysis by NE releases the fluorescent group, which can then be detected. As shown in Figure 6C, NE treated with empty vector *E. coli* W3110 supernatant successfully degrades the substrate, and fluorescence increased over time, indicating NE is active. By contrast, NE activity is strongly inhibited by addition of commercial eglin C or eglin C PS supernatant. The fluorescence signal from NE treated with eglin C NS supernatant increased over time, indicating NE remained active. However, the rate was slightly lower than that of empty vector NE control, suggesting that a small amount of eglin C is present in NS supernatant. This may be due to a small amount of eglin C released by cell lysis and detected by this sensitive assay. Overall, this result indicates that eglin C is successfully secreted via the MccV system and retains its activity.

Lastly, we tested the more complex polypeptide, human epidermal growth factor (EGF) that stimulates growth of epithelial cells through binding to a specific tyrosine kinase receptor (33). Mature EGF has three disulfide bonds. However, EGF is commonly purified from bacteria and refolded, suggesting the primary sequence is sufficient to develop the mature fold under oxidative conditions (34). Prior to testing the bioactivity of EGF, we measured the relative amount of EGF in supernatants from *E. coli* W3110 grown in mammalian cell culture medium (Fig. 6D) by colorimetric ELISA (enzyme-linked immunosorbent assay). In addition to the EGF NS control, we also included PS supernatant containing random peptide G3P2 as a non-specific control. We had to dilute the EGF PS supernatant 100-fold to obtain a similar OD_450_ as 1 ng/mL of EGF standard, suggesting our supernatant contained ~100 ng/mL EGF. No signal was detected from EGF NS supernatant or G3P2 PS supernatant. We then assayed the ability of *E. coli*-secreted EGF to activate the EGF receptor (EGFR) in cell culture. Transfected CHO cells expressing HA-tagged EGFR were treated with purified EGF standard, EGF PS supernatant, G3P2 PS supernatant. Cell lysates were immunoblotted for the presence of EGFR, phosphorylated (activated) EGFR, and β-actin (loading control) (Fig. 6E). EGF standard and EGF PS supernatant increased EGFR phosphorylation to a similar level, while no change in EGFR phosphorylation was observed from the G3P2 PS (non-specific peptide). No signal for EGFR was observed in non-transfected CHO cells. These results indicate that EGF can be secreted by *E. coli* through the MccV system, obtain its active form, and activate its target human receptor.

To further characterize these secreted peptides, we purified eglin C and EGF for analysis since both are derived from non-microbial sources and thus represent peptides far removed from native bacterial secretion. Each peptide was constructed with a C-terminal strep-tag and secreted into 0.5 L of bacterial growth medium. Peptides were purified from the supernatant via affinity chromatography, and purity was assessed by SDS-PAGE (Fig. S4A). We observed a single band for each peptide near the expected molecular weight. When analyzed by mass spectrometry, a single, dominant signal was observed for each peptide (EGF_strep: 7370.34 Da, Eglin C_strep: 9244.74 Da) (Fig. S4B and C). Each signal matched the theoretical mass of the respective peptide after SP_MccV_ cleavage within < 1 Da. For EGF, this profile matches the mature disulfide bond-containing form. Minor mass signals were observed, but none matched the mass of the peptides containing additional SP_MccV_ sequence residues. This indicates that SP_MccV_-cleaved peptides are the dominant species in the supernatant. The yields were 0.57 mg/L (EGF_strep) and 1.11 mg/L (Eglin C_strep) per OD_600_ under these test conditions. These yields are consistent with the range for the MccV system (0.19–7.25 mg/L per OD_600_) that we estimated by dot blot (Fig. 5B).

### The MccV system is functional in related Gram-negative bacteria

We have shown that the MccV system can secrete heterologous peptides from *E. coli* strains W3110 and BL21(DE3). Microcin T1SSs are predicted to be broadly distributed across Gram-negative bacteria (35) suggesting the MccV system may retain its function in non-*E. coli* strains. To make the secretion system more amenable to a wider range of bacteria, we cloned it into the broad-host-range plasmid, pMMB67EH (Fig. 7A). This configuration allows for expression of the POI and CvaAB under IPTG-inducible control. For testing purposes, we used pediocin PA-1 as the cargo peptide to provide a simple ZOI read out. We transformed this plasmid and the negative control plasmid lacking CvaAB into *E. coli* Nissle 1917, *Salmonella enterica* Ty21a, and *Vibrio cholerae* CVD103-HgR. *E. coli* Nissle is a commonly used probiotic strain (36). *S. enterica* Ty21a and *V. cholerae* CVD103-HgR have been explored as vaccine strains of human pathogens and are shown to be safe for human use (37, 38). Each bacterial strain expressing SP_MccV_-pediocin and CvaAB was able to inhibit the growth of *L. monocytogenes* (Fig. 7B), indicating the MccV system is functional in a probiotic *E. coli* strain and at least two additional Gram-negative species capable of gut colonization.

**Figure 7.**
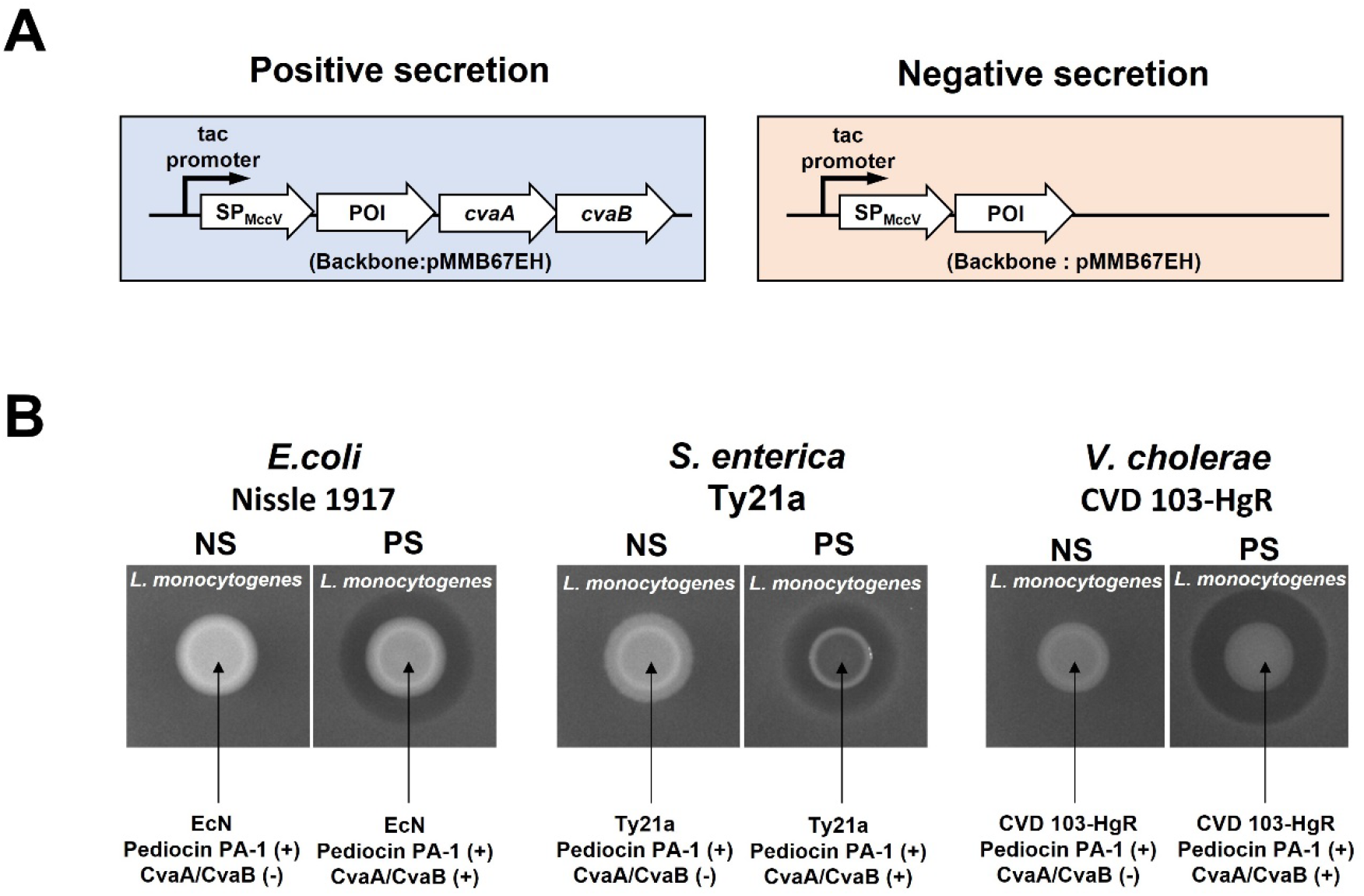
Compatibility of the MccV system with other Gram-negative bacteria. (A) One-plasmid peptide secretion system is shown. Peptide of interest (POI) conjugated with the MccV signal peptide (SP_MccV_) is expressed with CvaAB (positive secretion) or without CvaAB (negative secretion). (B) The result of pediocin PA-1 zone of inhibition assays is shown. Cultures of bacteria containing either pediocin PA-1 negative secretion (NS) or positive secretion (PS) spotted on a lawn of *L. monocytogenes* and 1 mM IPTG. The per species pair of NS and PS was spotted on the same agar plate and the spot images are prepared from the same plate picture, but separate plates were used for different bacterial species. The result is a representative of biological triplicate.

## Discussion

Secretion systems allow bacteria to influence their local environment through the release of a myriad of toxins, enzymes, signals, and colonization factors. Gram-positive bacteria need only secrete proteins across their cytoplasmic membranes to reach the external environment, while in Gram-negative bacteria, proteins must also transit the near-impenetrable outer membrane. T1SSs elegantly overcome this issue by providing a one-step transportation of proteinaceous cargo from the cytoplasm to the external environment, across both Gram-negative cell membranes. While factors affecting the secretion of large proteins, including toxins and enzymes, through T1SSs (1, 6, 7, 9, 10) have been studied, investigation of T1SSs that secrete microcins, small antibacterial peptides (< 10 kDa) (12, 13), has received considerably less attention, and no studies have been undertaken to understand features of the cargo peptide that influence their export.

Remarkably, we found that the MccV T1SS could secrete a wide range of synthetic and bioactive peptides with diverse sequence content. Despite the high glycine and hydrophobic residue content of native microcins, we found no strong association between peptide sequence content or chemistry and their secreted abundance. The only constraint on secretion through the MccV T1SS we could identify was peptide length. Peptides 26–66 amino acids were readily secreted, while peptides 116 amino acids long were not. The range of 66–116 amino acids straddles the length of the longest confirmed mature microcin (90 amino acids) (39), suggesting a cutoff that limits the natural size of microcins. This size cutoff may be explained by recent observations of the structure of a related PCAT, “PCAT1”, from the Gram-positive bacterium, *Clostridium thermocellum* (19, 20). An alternating-access model was proposed for its substrate transport through PCAT1. Once the signal peptide (SP) of the substrate is recognized by the flexible peptidase domain (PEP) of PCAT1, the translocating substrate would insert via its C-terminal region into the central cavity of the PCAT1 transmembrane domain. This cavity faces inward in the absence of ATP-binding. Following ATP-binding, PCAT1 changes from the inward to the outward cavity facing form. This enables the cargo in the PCAT1 transmembrane conduit that is captured by the PEP to be released into the extracellular space following cleavage of its SP. As a member of the PCAT family, CvaB may also follow this model. Substrates that are too long may not be efficiently loaded into the transmembrane conduit of CvaB or may hamper the conformational change to the outward form to translocate the substrate through the next parts of the secretion complex, CvaA and TolC. Another hypothesis is about TolC protein recruitment by the substrate. It is not clearly elucidated in the MccV T1SS, but in the hemolysin A (HlyA) T1SS, HlyA recruits TolC to assemble the HlyB (ABC transporter)-HlyD (membrane fusion protein)-TolC complex for its secretion (40). Considering that PhoA conjugated with the MccV SP was translocated to the inner membrane-facing periplasm (15), it may be possible that a peptide that is too long cannot recruit TolC properly, and remains stuck in the inner membrane-periplasm region. Other unexplored physicochemical properties, such as propensity to form second structures, may also influence peptide secretion and warrant further study.

The promiscuity of the MccV secretion systems opens opportunities for peptide research and delivery. We show that, in addition to antibacterial peptides like pediocin PA-1, the MccV system can secrete non-bacterial derived bioactive peptides as exemplified through secretion of a yeast pheromone (α-factor), a protease inhibitor (eglin C), and a human hormone (epidermal growth factor). A key benefit of the MccV system is the ease of use. Simply appending the MccV SP to the cargo peptide appears sufficient to drive the export of diverse peptides. As synthesis of long peptides remains chemically challenging, the MccV system could provide a facile method for small scale peptide production. An important consideration for this use is required peptide yield. Over 8 hours, *E. coli* W3110 secretes 0.19–7.25 mg/L of peptide per OD_600_. This yield is comparable with other Gram-negative secretion systems such as HlyA T1SS (nanobodies: 0.3– 1.3 mg/L) (10) or flagellar T3SS (cutinase 0.16 mg/L, human growth hormone: 0.15 mg/L) (39).

Another strategy (42) can produce higher peptide yields under fermentation (teriparatide: > 2 g/L) but requires cumbersome cell lysis, inclusion body recovery, and fusion protein cleavage to recover the desired peptide. We have not attempted to optimize export, but our studies using *E. coli* BL21(DE3) lacking proteases and lowering temperature indicate that peptide secretion abundance can be improved (Fig. S3). Investigations of several protein export systems have shown that secretion efficiency is highly dependent on the cargo, and solid rules for optimal secretion for any system remain elusive (1). Ideal conditions for peptide secretion through the MccV system likely have similar sequence dependency. In its current form, the MccV system may provide an opportunity for rapid small-scale analysis of novel peptides or peptide libraries.

As highlighted by other secretion systems (43–46), extracellular secretion via the MccV system could provide opportunities for *in vivo* peptide delivery. Oral delivery of peptides is challenging because they are quickly degraded during passage through the stomach (47). Many bacteria can survive this passage, so could be used to overcome this barrier and begin delivering bioactive peptides once in the gut to affect a wide range of outcomes, such as altering metabolism, influencing immune responses, or treating cancers. The MccV system is active in several bacteria that have tropisms for colonizing different gut locations which could be leveraged to deliver peptides to specific areas of the intestinal tract. Bacteria secreting peptides through the MccV system could act as on-site production factories to generate a constant localized flow of peptides for sustained effects. The use of bacteria like *Salmonella* may even offer the chance for intracellular delivery. Bacteria that colonize other areas of our body could be used similarly for niche-specific delivery.

## Materials and Methods

Bacterial and yeast strains used in this study are listed in Table S3. Standard techniques in molecular cloning were used to construct plasmids (48). “Peptides” R package (49) was used to calculate peptide chemical characteristics. Western and dot blots were performed using standard techniques for immunoassay. Activity of peptides was assayed based on the native biological function. Peptides were purified using Strep-Tactin XT Sepharose and mass analyzed using a Thermo Orbitrap Fusion Tribrid mass spectrometer. The experimental procedures are detailed in Supporting Information Appendix: Materials and Methods.

## Acknowledgments

*S. cerevisiae* CMY 740-1D was acquired from the Matouschek lab at The University of Texas at Austin. We thank Dr. Chris Yellman for technical discussion. Plasmids for amplification of the MccV system(pHK22) were kindly provided by the Kolter lab at the Harvard Medical School. This work was supported by DARPA (HR0011-19-2-001 to BWD), NIH (R01 AI125337, R01 AI148419, and R21 AI159203 to B.W.D.;), DTRA (HDTRA1-17-C-0008 to BWD), Tito’s Handmade Vodka (to B.W.D.), CPRIT (RR160023 to DJL).

