## Supporting Information for "Export of diverse and bioactive peptides through a type I secretion system"

**Author Contributions:** S-Y.K., J.K.P., M.G-M., M.S.T., D.J.L., and B.W.D. designed research; S-Y.K., M.G-M., and M.S.T. performed research; S-Y.K., J.K.P., M.G-M., M.S.T., D.J.L., and B.W.D. analyzed data; and S-Y.K., J.K.P., M.G-M., D.J.L., and B.W.D. wrote the paper.

**Competing Interest Statement:** No, the authors declare no competing interest.

**Keywords:** Secretion, Peptide, Gram-negative bacteria, T1SS

**This PDF file includes:**

Supporting text
Supporting References
Figures S1 to S4
Tables S1 to S5

### Supporting text

### Supplementary Materials and Methods

#### Bacterial and yeast strains, growth conditions and genetic modification

Bacterial and yeast strains used in this study are listed in Table S3. For plasmid construction and molecular cloning, *E. coli* DH5α competent cells (New England Biolabs, Cat# C2987H) were used. All Gram-negative bacteria were grown in lysogeny broth (LB) media at 37 °C with shaking at 220 rpm, unless otherwise stated. *L. monocytogenes* was grown in tryptic soy broth (TSB) media at 37 °C with shaking at 220 rpm. *S. cerevisiae* was grown in yeast peptone dextrose (YPD) media at 30 °C with shaking at 275 rpm. As required, the following antibiotics were used: carbenicillin 75 µg/mL, kanamycin 50 µg/mL, streptomycin 100 µg/mL, and chloramphenicol 10 µg/mL.

#### Construction of plasmids

Standard techniques in molecular cloning were used to construct plasmids (1). All plasmids, primers, and gBlocks™ used in this study are listed in Tables S4 and S5. Primers and gBlocks™ were ordered from Integrated DNA Technologies (IDT). Random synthetic peptide inserts for cloning were built by using either reduced random codon (NNK) containing primer sets, or gBlocks™ and cognate primer sets. The gBlocks™ contain nucleotide sequences that are reverse-translated with codon optimization from peptide sequences randomly generated by (https://www.genscript.com/sms2/index.html). Other inserts containing peptides of interest were constructed by using either codon optimized open reading frame (ORF)-containing gBlocks™ or primers, cognate primer sets, and the selected plasmids mentioned below.

Two-plasmid secretion system: Plasmids pBAD18, or its derivative pSK00, which encodes MccV SP, were used to express peptides of interest. Each peptide of interest was conjugated with the MccV signal peptide (CvaC15, MRTLTLNELDSVSGG) at the N-terminus and cloned into pBAD18. If required, the V5 tag with two additional N-terminal glycine residues (GGGKPIPNPLLGLDST) was conjugated at the C-terminus. We used pHK22 (2) as a template to amplify *cvaA* and *cvaB* genes. CvaAB were cloned into pACYC184 for constitutive expression. We refer to this plasmid as pSK01. We also constructed pSK02, by introducing a point mutation in *cvaB* of pSK01 to express mutant-type CvaB (C32S).

One-plasmid secretion system: A broad-host-range vector, pMMB67EH was used to construct pSK03. Amplified *cvaA* and *cvaB* as described above were cloned into pMMB67EH. Pediocin PA-1 is cloned into pSK03 using primers and gBlocks™ described in Table S4.

#### Calculating peptide properties

We used the "Peptides" R package (3) to calculate charge and hydrophobicity of the random synthetic peptides, as described in the Peptides package documentation. Using the same package, the amino acid frequencies in a particular class were determined per group and normalized by the total number of amino acids in the group (# of certain amino acids/# of total amino acids) to obtain percent composition.

**Zone of inhibition assay**

To observe the zone of inhibition (ZOI) by MccV or MccV_V5, *E. coli* K-12 W3110 wild-type was used as the susceptible strain. Overnight culture of the susceptible strain was diluted to $OD_{600}$ = 0.001 in LB agar medium with 0.2% (v/v) arabinose and solidified. Overnight cultures of the secreting strains, including empty vector, positive secretion, negative secretion, and protease-deficient secretion (Fig. 1) strains, were centrifuged at 5,000×*g* for 5 min and the pellets were re-suspended in 100 µl of fresh LB medium. 5 µl of the suspensions were spotted on the solidified agar and pictures were taken after overnight incubation at 37 °C.

Similarly, *L. monocytogenes* was used as a susceptible strain to detect ZOI by pediocin PA-1. Overnight *L. monocytogenes* culture diluted to $OD_{600}$ = 0.001 in TSB agar with 0.2% (v/v) arabinose (Fig. 6A) or 1 mM of IPTG (Fig. 7B) and solidified. The overnight cultures of *E. coli* W3110 including empty vector, positive secretion, negative secretion, and protease-deficient secretion (Fig. 6A) and *E. coli* Nissle, *S. enterica*, and *V. cholerae* containing either positive or negative secretion (Fig. 7B) were enriched and spotted in a same way above. The pictures were taken after overnight incubation at 37 °C.

*S. cerevisiae* CMY 740-1D was used as a susceptible strain to detect ZOI caused by secreted α-factor. An overnight culture was diluted to $OD_{600}$ = 0.01 in yeast peptone glycerol (YPG) agar medium with 0.2% (w/v) arabinose and solidified. Overnight cultures of secreting strains (*E. coli* W3110) were enriched as described above and 5 µl were spotted on the solidified YPG agar plate. Pictures were taken after 30 hours incubation at 30 °C.

**Western blot**

To detect MccV_V5 (Fig. 2), an overnight *E. coli* W3110 liquid culture was diluted in LB medium to $OD_{600}$ = 0.5. The culture was induced with 0.2% (v/v) arabinose for 2 hours and normalized to $OD_{600}$ = 1.0. 250 µl of culture was taken and centrifuged at 5,000×*g* for 5 min to separate the supernatant and cell pellet. The Invitrogen Novex Tricine Gel System (Thermo Fisher Scientific) was used to perform SDS-PAGE. The supernatant and cell pellet were each suspended in Tricine SDS Sample Buffer (Cat# LC1676) with Sample Reducing Agent (Cat# NP0009), and 10 µl was loaded into each well of 16% Tricine gel (Cat# EC66952) with 10 µl of SeeBlue Plus2 Pre-stained Protein Standard (Cat# LC5925) and migrated with Tricine SDS Running Buffer (Cat# LC1675). Then, the peptides were transferred to a nitrocellulose membrane

(Cat# LC2000). The membrane was blocked with 5% (w/v) low-fat milk in TTBS (50 mM Tris-HCl, pH 7.5, 150 mM NaCl, 0.05% Tween-20 (v/v)) and proteins were labeled by incubation with the selected primary antibody, anti-V5 antibody (Sigma-Aldrich Cat# V8012) or anti-DnaK antibody (Enzo Cat# ADI-SPA-880) 1:5000 diluted in 1% BSA (Bovine Serum Albumin) in TTBS. LI-COR IRDye 800CW Goat Anti-Mouse IgG (Li-cor Biosciences, Cat# 926-32210) was used as the secondary antibody 1:5000 diluted in 5% (w/v) low-fat milk in TTBS. The Li-Cor Odyssey Clx Near IR imaging system was used for visualizing. Band intensities were measured using Image Studio software (https://www.licor.com/bio/image-studio-lite/download).

**Dot blot**

*E. coli* culture samples for dot blot were grown either using test tubes, following the growth conditions mentioned for western blot above, or in a 96-well deep well plate (NEST Scientific Cat# 503501). When using a deep well plate, a single colony was inoculated into 1 ml of LB medium in each well and incubated at 37 °C with shaking at 1,000 rpm. The plate was sealed by a permeable membrane (Diversified Biotech Cat# BEM-1) for proper air circulation. After overnight growth, cultures were diluted in 500 µl of fresh LB medium to $OD_{600}$ = 0.5 with 0.2% (w/v) arabinose and incubated at 37 °C with shaking at 1,000 rpm for 8 hours. The plate was centrifuged at 4,000 rpm for 10 min to collect supernatants samples. A nitrocellulose membrane (GE Healthcare Life Sciences, Cat# 10600010) was inserted into a 96-well Bio-Dot Microfiltration Apparatus (Bio-Rad, Cat# 1706545), and 100 µl of each supernatant was loaded onto a well in the apparatus and filtered following the manufacturer’s protocol. To perform dot blots for total cell lysate samples, induced cultures were boiled for 20 mins and centrifuged at 5,000×*g* for 5 min to separate supernatant and cell debris. 100 µl of the collected supernatants were loaded onto wells. After samples passed through the membrane, the membrane was washed twice with TTBS and removed from the apparatus. Blocking, antibody incubation, visualization, and signal intensity calculations were done as described in the western blot section. The raw dot blot signal intensity of the supernatant sample was divided by the $OD_{600}$ value of the original culture. Then, the normalized intensity of negative secretion (no CvaAB) was subtracted from the normalized positive secretion intensity. The obtained values were referred to as “relative secretion level” in Figure 3 and Figure 4. Raw dot blot signal intensity of total cell lysate sample was divided by $OD_{600}$ value of culture. Next, normalized intensity of total cell lysate was subtracted from intensity of empty vector culture. The obtained values were referred to “relative cellular abundance level” in Figure 3 and Figure 4.

**Secreted peptide quantification**

Positive and negative secretion supernatants of G1P9, G2P9, G3P2 and G4P7 (Fig. 5) were diluted in fresh LB, and 100 µl of the diluents were dot blotted. Supernatant samples were diluted accordingly to be in the range of the standard curve. Signal intensity was normalized as described above. Secreted peptide concertation was determined by comparison to a standard curve (signal intensity vs. μM). The standard peptide (SFRNGVGSGAKKTSFRRAKQGGKPIPNPLLGLDST) was synthesized by GenScript with ≥90% purity. The secreted peptide concentrations were converted to mg/L per $OD_{600}$.

**Elastase inhibition assay**

Eglin C positive secretion, negative secretion and empty vector containing *E. coli* W3110 were grown overnight, and 100-fold diluted in fresh LB medium. Once the cultures reached $OD_{600}$ = 0.5, the medium was replaced with M9 minimal medium supplemented with 0.4% (v/v) glycerol and 0.2% (w/v) arabinose and grown overnight for induction. Cell-free supernatants were collected by centrifuging the cultures at 5,000×*g* for 5 min and filtering them using a PES 0.22 μm filter membrane (Celltreat Scientific, Cat# 229747). The obtained samples' protease inhibition activity was tested using a Neutrophil elastase inhibitor screening kit (Abcam, Cat# ab118971) following the manufacturer's protocol. For a positive control, N-Acetyl-eglin C peptide was purchased from Enzo Life Sciences (Cat # ALX-201-006-MC01) and diluted at 1.25 μM in empty vector supernatant prepared as described above.

**ELISA**

EGF positive secretion, negative secretion, and G3P2 positive secretion *E. coli* W3110 were grown overnight, and 100-fold diluted in fresh LB medium. Once the cultures reached $OD_{600}$ = 0.5, the medium was replaced with Ham's F-12 medium (Thermo Fisher Scientific, Cat# 11765070) supplemented with 1 mg/ml BSA and 0.2% (w/v) arabinose and grown overnight for induction. Cell-free supernatants were collected as described, and the supernatants were directly used to detect secreted EGF by using Human EGF ELISA kit (Boster Bio, Cat# EK0325) following the manufacturer's protocol. Standard EGF provided by the company was used as a positive control.

**Mammalian cell culture, transfection, and EGFR phosphorylation assay**

CHO-K1 cells (ATCC, Cat# CCL-61) were grown in six-well plates at $1 \times 10^{6}$ cells/well and transfected with 1.5 μg of pcDNA-EGFR-HAtag DNA using PEI (polyethyleneimine, Fisher Scientific, Cat# NC1014320) as described before (4) in a ratio of 3:1. After 18 hours, cells were washed three times with 2 ml Ham's F-12 supplemented with 1 mg/ml BSA and incubated in this medium for 3 hours at 37 °C to serum starve. The different ligands were added in specific wells for 5 min: a control of 100 ng/ml EGF purified as described by Qiu *et al*. (5), the same supernatant samples used for EGF ELISA assay with estimated concentration of 100 ng/μl, and a dilution of 1:100 in Ham's media for 5 min at 37 °C. A G3P2 supernatant sample was used as a non-specific peptide for a negative control. Wells were washed with ice-cold phosphate-buffered saline and then lysed for 30 min at 4 °C in 250 µl of RIPA buffer supplemented with 1 mM activated sodium orthovanadate, a Pierce protease inhibitor minitablet (Thermo Fisher Scientific, Cat# A32955), and Benzonase nuclease (Sigma-Aldrich, Cat# E1014). Total protein concentrations of clarified lysates were determined using the BCA (Bicinchoninic Acid) assay and lysates were normalized to the lowest total protein content using RIPA buffer. Normalized amount of protein lysates was mixed with sample buffer and boiled, separated by SDS-PAGE 4–12%, and transferred onto a nitrocellulose membrane. The membrane was blocked with 3% (w/v) low-fat milk in TBS, and proteins were detected by incubation with: rabbit anti-EGF Receptor (D38B1) (Cell Signaling, Cat# 4267), rabbit anti-phospho-EGFR pTyr1068 antibody (Thermo Fisher Scientific, Cat # 44-788G), rabbit anti-β-Actin (Cell Signaling Technology, Cat # 4968), and the secondary antibody Goat anti-rabbit-680RD (Li-Cor, Cat # 926-68071). Visualization is done as described in the western blot section.

**Recombinant peptide purification**

We constructed two pBAD18 derivative plasmids that express recombinant EGF (epidermal growth factor) and Eglin C, respectively. Each recombinant peptide was conjugated with the MccV signal peptide (CvaC15, MRTLTLNELDSVSGG) at the N-terminus, and Strep-tag® II with two glycine residues (GGWSHPQFEK) at the C-terminus. The plasmids were transformed into *E. coli* BL21(DE3) containing pSK01, which constitutively expresses CvaAB, to generate EGF_strep and Eglin C_strep positive secretion strains. Overnight EGF_strep and Eglin C_strep positive secretion cultures were diluted into 0.5 L of fresh M9 minimal media at final $OD_{600}$ = 0.5, with 0.2% (v/v) glycerol and 0.1% (w/v) casamino acids. The cultures were induced with 0.2% (w/v) of arabinose overnight at 30 °C. Supernatants were collected by centrifuging the cultures at 6,000×g for 20 min, then filtered through PES 0.22 µm filter membranes (Genesee Scientific, Cat# 25-233). 250 µL bed volumes of Strep-Tactin® XT Sepharose™ (Cytiva, Cat# 29401324) was equilibrated with PBS. Each recombinant peptide in PBS-equilibrated supernatant was bound to the resin and washed once with PBS buffer. Each peptide was eluted using 2 mL of elution buffer (PBS containing 50 mM biotin). Each peptide solution was concentrated by a 3 kDa cut-off ultra-filtration (MilliporeSigma Cat# UFC500324). Purified peptides were resolved by SDS-PAGE and stained with SimplyBlue™ SafeStain (Thermo Fisher Scientific, Cat# LC6065).

**Mass spectrometry**

Purified peptide solutions prepared above were desalted using C18 HyperSep™ SpinTip Microscale SPE Extraction Tips (Thermo Fisher Scientific, Cat# 60109-412) and eluted into buffer (60% (v/v) acetonitrile, 1% (v/v) formic acid, and 0.05% (v/v) trifluoroacetic acid). The peptide solutions were directly analyzed by mass spectrometry without further treatments. The mass spectrum of each sample was determined by the UT Austin Center for Biomedical Research Support Biological Mass Spectrometry Facility (RRID:SCR_021728) using a Thermo Orbitrap Fusion Tribrid mass spectrometer. Theoretical masses of peptides were calculated from https://www.peptidesynthetics.co.uk/tools/.

**Peptide quantitative assay**

The concentrated peptide solutions prepared as described in “Recombinant peptide purification and SDS-PAGE” were quantified using Pierce™ Quantitative Peptide Assays & Standards (Thermo Fisher Scientific, Cat# 23290) as per manufacture’s protocol.

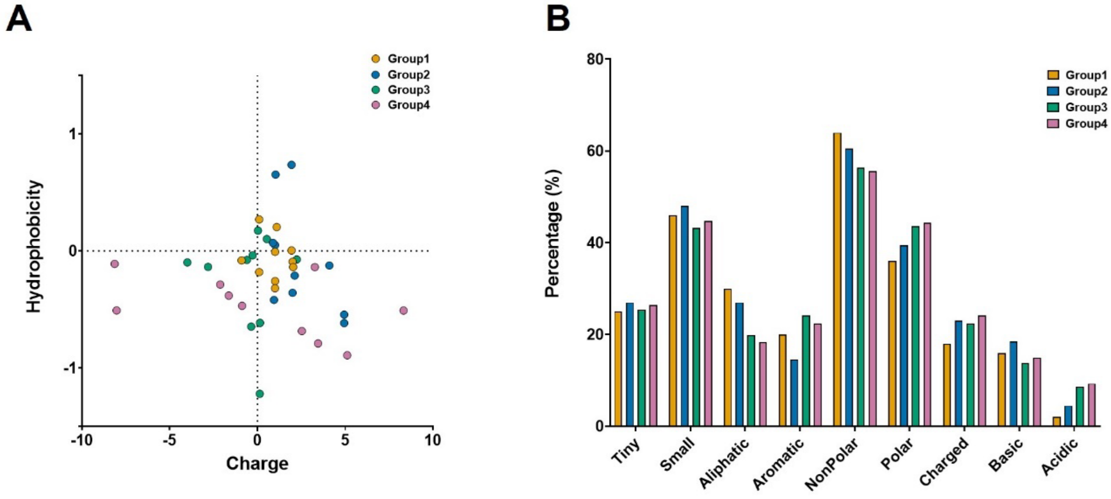


**Fig. S1. Properties of random synthetic peptides**. (A) Theoretical charge (at pH = 7.0) versus hydrophobicity of peptides is shown as a scatter plot. Different groups are represented as a different color; Group 1 as orange, Group 2 as blue, Group 3 as green, and Group 4 as purple. (B) The composition of amino acids belonging to a particular class was calculated per group. Amino acids were classified as “Tiny” (alanine, cysteine, glycine, serine, threonine), “Small” (alanine, cysteine, aspartic acid, glycine, asparagine, proline, serine, threonine, valine), “Aliphatic” (alanine, leucine, valine), “Aromatic” (phenylalanine, histidine, tryptophan, tyrosine), “Non-Polar” (alanine, cysteine phenylalanine, glycine, leucine, methionine, proline, valine, tryptophan, tyrosine), “Polar” (aspartic acid, glutamic acid, histidine, lysine, asparagine, glutamine, arginine, serine, threonine), “Charged” (aspartic acid, glutamic acid, histidine, lysine, arginine), “Basic” (histidine, lysine, arginine), and “Acidic” (aspartic acid, glutamic acid). Group colors are the same as in (A).

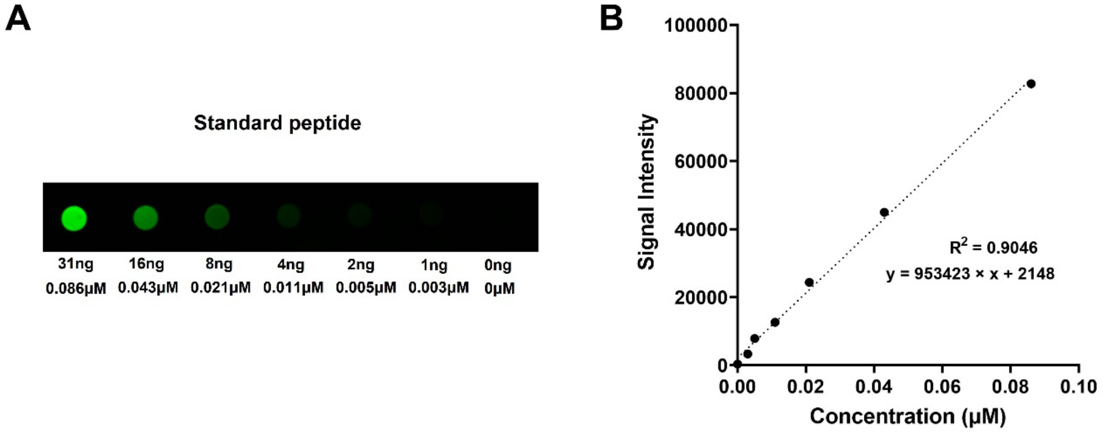


**Fig. S2. Standard curve of V5-tagged synthetic peptide.** (A) Result of dot blot for V5-tagged standard peptide is shown. Standard peptide was loaded with serial 2-fold dilution, and the total amount (ng) and concentration (μM) of the peptide in each well are shown. A representative dot blot image prepared from a single membrane is shown. (B) The mean of four signal intensity values vs. concentration (μM) of standard peptide is plotted. Simple linear regression assay was performed. The best-fit slope is shown as a line with R-squared value and standard curve equation.

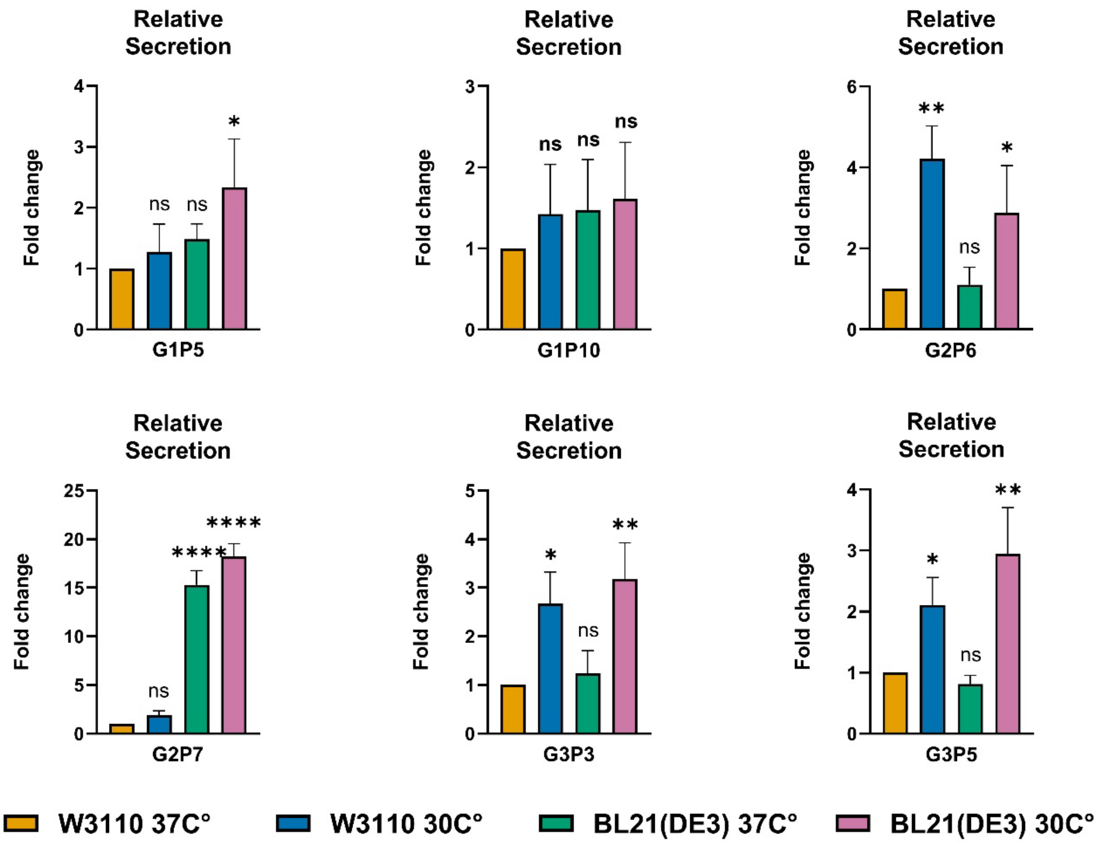


**Fig. S3. Optimization of secretion level of random peptides.** Relative secretion levels and cellular abundance levels of random peptides (G1P5, G1P10, G2P6, G2P7, G3P3 and G3P5) produced from *E. coli* W3110 or BL21(DE3) strains grown at 37 °C or 30 °C (represented as W3110 37 °C: orange, W3110 30 °C: blue, BL21(DE3) 37 °C: green, and BL21(DE3) 30 °C: purple). Peptide levels were normalized to our standard condition (W3110 37 °C), and the results are shown as fold change. Relative secretion levels and cellular abundance levels were calculated as described in “Material and Methods: Dot blot”. The mean of biological triplicate is shown with standard deviation. Adjusted *P*-values were calculated by ANOVA with Dunnett’s multiple comparison test (vs. W3110 37 °C) and are shown as not significant (ns = $P > 0.05$) or the number of asterisks to indicate significance level (* = $P < 0.05$, ** = $P < 0.01$, **** = $P < 0.0001$).

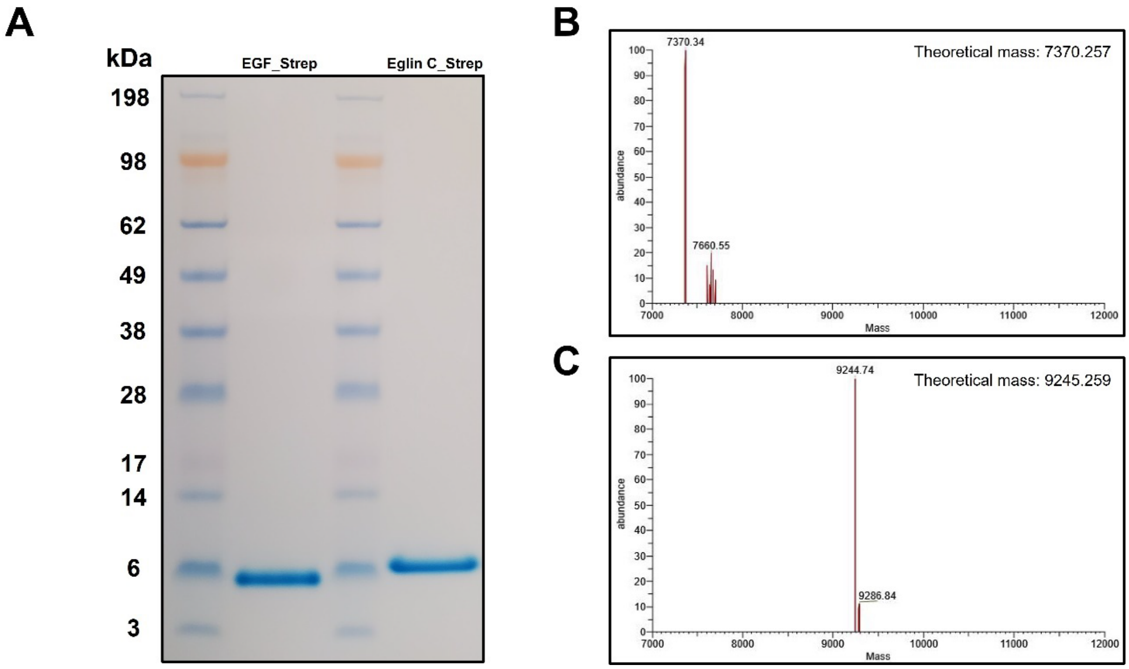


**Fig. S4. Purification of recombinant peptides from supernatant.** (A) The result of SDS-PAGE of strep-tagged EGF (EGF_strep) and Eglin C (Eglin_C strep) is shown. Molecular weights (kDa) of protein ladders are presented on the left. (B and C) Deconvoluted mass spectra of (B) EGF_strep, or (C) Eglin C_strep solution is shown. Average theoretical masses of (B) EGF_strep and (C) Eglin C_strep without the MccV signal peptide are also shown. Cysteine oxidation is considered for the EGF_strep theoretical mass calculation. The y-axis represents the relative abundance (%) calculated from the relative intensity of the detected mass, and the x-axis represents the mass (Da). Averages of deconvoluted masses are shown above the respective peaks.

**Table S1. Properties of random synthetic peptides.**

| ID | Group | Sequence | Length (aa) | MW | GRAVY | Charge |
|---|---|---|---|---|---|---|
| G1P1 | 1 | GIGWMLSRARGGGKPIPNPLLGLDST | 26 | 2664.15 | -0.09 | 2 |
| G1P2 | 1 | IVHRYPRICYGGGKPIPNPLLGLDST | 26 | 2837.37 | -0.14 | 2.03 |
| G1P3 | 1 | SHSMVSVLVRGGGKPIPNPLLGLDST | 26 | 2632.1 | 0.2 | 1.09 |
| G1P4 | 1 | GIYGHIVVYWGGGKPIPNPLLGLDST | 26 | 2724.18 | 0.27 | 0.09 |
| G1P5 | 1 | SGLRPWMASVGGGKPIPNPLLGLDST | 26 | 2621.08 | -0.01 | 1 |
| G1P6 | 1 | LSMSICMRPKGGGKPIPNPLLGLDST | 26 | 2683.28 | 0 | 1.94 |
| G1P7 | 1 | VNDRLKLQWVGGGKPIPNPLLGLDST | 26 | 2788.27 | -0.26 | 1 |
| G1P8 | 1 | PGLIDVSYWHGGGKPIPNPLLGLDST | 26 | 2704.1 | -0.08 | -0.91 |
| G1P9 | 1 | SHSVAPWSLQGGGKPIPNPLLGLDST | 26 | 2628.99 | -0.18 | 0.09 |
| G1P10 | 1 | RYFTLNNFGWGGGKPIPNPLLGLDST | 26 | 2835.24 | -0.32 | 1 |
| G2P1 | 2 | GSVTRFISFWHMMLLCGMLVGGGKPIPNPLLGLDST | 36 | 3846.7 | 0.65 | 1.03 |
| G2P2 | 2 | SVIRINCLVLVGRLVLGTQVGGGKPIPNPLLGLDST | 36 | 3670.46 | 0.74 | 1.94 |
| G2P3 | 2 | GLWRAFMPWWSFFGVVRDSGGGGKPIPNPLLGLDST | 36 | 3919.56 | 0.05 | 1 |
| G2P4 | 2 | ALLRRDVSFRFWHRSVLYVRGGGKPIPNPLLGLDST | 36 | 4094.82 | -0.13 | 4.09 |
| G2P5 | 2 | NCDRWRGRWVRFILWFGRGKGGGKPIPNPLLGLDST | 36 | 4126.85 | -0.54 | 4.94 |
| G2P6 | 2 | HNPRRHWMGLITLKLMSCDLGGGKPIPNPLLGLDST | 36 | 3939.72 | -0.21 | 2.12 |
| G2P7 | 2 | SVCPAFRVDTFRSTGDKYSNGGGKPIPNPLLGLDST | 36 | 3768.26 | -0.42 | 0.94 |
| G2P8 | 2 | RQRDRCNPWMPIRAKRSLLVGGGKPIPNPLLGLDST | 36 | 4013.79 | -0.62 | 4.94 |

| | | | | | | |
|---|---|---|---|---|---|---|
| G2P9 | 2 | GNARCSMCEIRPLMVWTVSTGGGKPIPNPLLGLDST | 36 | 3772.48 | 0.07 | 0.88 |
| G2P10 | 2 | GVSYNTMVQRRGGDPARALMGGGKPIPNPLLGLDST | 36 | 3697.29 | -0.36 | 2 |
| G3P1 | 3 | LSNGQCNMHCCPCLEYQDYHKHYSNTESFKQLVWMTHICDNYALSHRAKWGGGKPIPNPLLGLDST | 66 | 7553.66 | -0.62 | 0.14 |
| G3P2 | 3 | HGVQGINIEKQPKRNNPENEQTRMKMRQERDWSCFAMFHAITRDDSYEQNGGGKPIPNPLLGLDST | 66 | 7583.53 | -1.22 | 0.13 |
| G3P3 | 3 | ETCHWMELHIPLFETDSFKPYDPKSLDSGHCLYYGFFFKYIGGLHAMCMYGGGKPIPNPLLGLDST | 66 | 7488.72 | -0.14 | -2.82 |
| G3P4 | 3 | VQMIWGFCSGWPMITWYAMMFAHIQWAFWNTKISSRGWEFMASAWPEYFVGGGKPIPNPLLGLDST | 66 | 7633 | 0.17 | 0.03 |
| G3P5 | 3 | IMECSTLICTTLDCFICVQGQWRYCMWNQCWCVCMNWTKTNCAQYSVAKHGGGKPIPNPLLGLDST | 66 | 7466.9 | 0.1 | 0.53 |
| G3P6 | 3 | IICWWPHPQCCWNFEYCFRKNYLTCFQCSEQYSTTFAPVNAFPTIWQIIYGGGKPIPNPLLGLDST | 66 | 7710 | -0.04 | -0.28 |
| G3P7 | 3 | GSVIVLDWVQGTNLVHKQHNTINRRHHHSHQEMYPWPTVFEMHGNRIIEEGGGKPIPNPLLGLDST | 66 | 7527.57 | -0.65 | -0.36 |
| G3P8 | 3 | HVPKWWYKGFDWTTQVWPYAAMLGFINAHHLDTVMIKIHLGAFHNCDWVEGGGKPIPNPLLGLDST | 66 | 7529.77 | -0.08 | -0.61 |
| G3P9 | 3 | VTTFKLWAKALVAFMYDADHHVNDFLPTLYRVYTTMNIWHFKHKCPCTSYGGGKPIPNPLLGLDST | 66 | 7543.86 | -0.07 | 2.23 |
| G3P10 | 3 | DDDDELLRLCDNITFFMMCIHEFTMKPWFKTIWFLMCWNAFLNQGSNSTHGGGKPIPNPLLGLDST | 66 | 7625.89 | -0.1 | -4 |
| G4P1 | 4 | YMCWHQAINVMEPCAQFYQDIVLSSRVQWQDMDMSMPRLYKMQYVAKSHF | 116 | 13715.07 | -0.29 | -2.13 |

| | | | | | | |
|---|---|---|---|---|---|---|
| | | SIMYFIHREDIQSSGCCDCNCNPVRKI FCTRYVEMVDCGYWRHFWLPPEEG GGKPIPNPLLGLDST | | | | |
| G4P2 | 4 | FIIYIPKVYDWYAASMGCTYPDGLGFR MVSMRVIWYAWYVYSCYTAVETNKF GDCGCGNYNHPDQEQIRYKTSEQES TKMFFRWAPEWQNNHRHHLMVEMG GGKPIPNPLLGLDST | 116 | 13608.64 | -0.47 | -0.88 |
| G4P3 | 4 | HSYCEWLMAKILSMGEQWWHKYYFG LRHQFNVSKGYHFNTSFDFHRCGEP KNNYYARTHCEKEYSNDDVHPRQNS MGRWGAEWPALIFKPLFGINKWGNF GGGKPIPNPLLGLDST | 116 | 13688.57 | -0.79 | 3.45 |
| G4P4 | 4 | SGIYCQITRWVHPFSESTQNMDNMA NTKRKPQWHYPRRHQHKEAQFVFFG IARGWPFQFFEQGWVTIDVHEEHLWI FCFGERNMEHGNVERAPTISSRIKGG GKPIPNPLLGLDST | 116 | 13624.55 | -0.69 | 2.52 |
| G4P5 | 4 | KHWKQRHVYYRQAVYQQQKCQMYN VPYSPASGCFNCQPNCHRKDFDWRD HTGYCFKMLQFHNFITPKRCTESALF AGQECRPDFAQESGAQKDFSGMHP GGGGKPIPNPLLGLDST | 116 | 13393.27 | -0.89 | 5.11 |
| G4P6 | 4 | FNPESSHQPSTIPKSHFRIICHFVRDW HFPCGSWTFSVVDIYFCEMYTTLGNP HGFVICCTYGSQYSGDNRCADKLER HPAMMENTYGWHGHTSAGLAQPGG GKPIPNPLLGLDST | 116 | 12962.73 | -0.38 | -1.64 |
| G4P7 | 4 | DKMEPQWNHSPRCFASLCCGGSHT MMAWNHVISWKGRDLVIGVNRHCAT PPHSQFWHNAWWQGFIKHVIETPRL GNMAKNMQCFAACALAVAFPVDISQL GGGKPIPNPLLGLDST | 116 | 12878.07 | -0.14 | 3.27 |
| G4P8 | 4 | RGKWMMCNTHMAHWNTYFWDGSA VHLTDDFYRNGPAKSYNLFMVQNHK ESRHNYKVCFFCYLITTYITIAKHKRM NENYWWMAQVYLKFVRWHARNCCY AGGGKPIPNPLLGLDST | 116 | 13910.21 | -0.51 | 8.32 |

| | | | | | | |
|---|---|---|---|---|---|---|
| G4P9 | 4 | VMQGDTWWECEPSEAEIQMLYWPW<br>GSQKDPIDWAYLCDTWKYTGDLCSG<br>GPEQPDEHRIHDAIGRAFYRPCPSLN<br>MYYLSQRWAIFDTHNSLAAGSYCFM<br>GGGKPIPNPLLGLDST | 116 | 13255.01 | -0.51 | -8.03 |
| G4P10 | 4 | TGRQVTEVTVWHALTTICGISELEFTY<br>GACPMWENMELEKFSGNVCYELQDH<br>CFCDWWQYTERCLENLPMIELPIQWK<br>PFTLHEWWIIGRCPLTIMNSWAGGGK<br>PIPNPLLGLDST | 116 | 13468.68 | -0.11 | -8.15 |

* aa: amino acids, GRAVY: Grand average of hydropathy, MW: Molecular weight (g/mol).

**Table S2. Selected bioactive peptides.**

| Name | Function | Origin | Length (aa) | MW |
|---|---|---|---|---|
| Pediocin PA-1 | Anti-bacterial peptide, strongly inhibits *L. monocytogenes* | *Pediococcus acidilactici* | 44 | 4628.19 |
| α-factor | Peptide pheromone, arrests cell cycle | *Saccharomyces cerevisiae* | 13 | 1683.99 |
| Eglin C | Protease inhibitor, inhibits neutrophil elastase | *Hirudo medicinalis* | 70 | 8091.05 |
| EGF | Epidermal growth factor, activates EGFR signaling | *Homo sapiens* | 53 | 6353.21 |

* aa: amino acids, MW: Molecular weight (g/mol).

**Table S3. Bacterial and yeast strains.**

| Name | Description | Source |
|---|---|---|
| *Escherichia coli* DH5α | *fhuA2 lac(del)U169 phoA glnV44 Φ80' lacZ(del)M15 gyrA96 recA1 relA1 endA1 thi-1 hsdR17* | NEB® |
| *Escherichia coli* W3110 | Wild-type | Lab stock |
| *Escherichia coli* BL21(DE3) | *fhuA2 [lon] ompT gal (λ DE3) [dcm] ∆hsdS*<br>*λ DE3 = λ sBamHIo ∆EcoRI-B int::(lacI::PlacUV5::T7 gene1) i21 ∆nin5* | NEB® |
| *Escherichia coli* SM10 (λpir) | *thi thr leu tonA lacY supE recA*::RP4-2-Tc::Mu Km *λpir* | Lab stock |
| *Escherichia coli* Nissle 1917 | Wild-type, Probiotic strain | Lab stock |
| *Salmonella enterica* Ty21a | CDC 2861-79 | ATCC® 33459 |
| *Vibrio cholerae* CVD103-HgR | mutant from clinical isolate 569B | ATCC® 55456 |
| *Listeria monocytogenes* EGD-e | Wild-type | Lab stock |
| *Sacharomyces cerevisea* CMY 740-1D | *MATa his3Δ1 leu2-3,112 trp1-289 ura3-52 bar1::loxP* | Matouschek Lab |
| SK00 | *E. coli* W3110, pBAD18-Km/pACYC184 | This study |
| SK01 | *E. coli* W3110, MccV positive secretion, encoding pSKP00/pSK01 | This study |
| SK02 | *E. coli* W3110, MccV negative secretion, encoding pSKP00/pACYC184 | This study |
| SK03 | *E. coli* W3110, MccV protease-deficient secretion, encoding pSKP00/pSK02 | This study |
| SK04 | *E. coli* W3110, MccV_V5 positive secretion, encoding pSKP01/pSK01 | This study |
| SK05 | *E. coli* W3110, MccV_V5 negative secretion, encoding pSKP01/pACYC184 | This study |
| SK06 | *E. coli* W3110, MccV_V5 protease-deficient secretion, encoding pSKP01/pSK02 | This study |
| SK07 | *E. coli* W3110, G1P1 positive secretion, encoding pSKP02/pSK01 | This study |
| SK08 | *E. coli* W3110, G1P1 negative secretion, encoding pSKP02/pACYC184 | This study |
| SK09 | *E. coli* W3110, G1P2 positive secretion, encoding pSKP03/pSK01 | This study |
| SK10 | *E. coli* W3110, G1P2 negative secretion, encoding pSKP03/pACYC184 | This study |
| SK11 | *E. coli* W3110, G1P3 positive secretion, encoding pSKP04/pSK01 | This study |
| SK12 | *E. coli* W3110, G1P3 negative secretion, encoding pSKP04/pACYC184 | This study |
| SK13 | *E. coli* W3110, G1P4 positive secretion, encoding pSKP05/pSK01 | This study |
| SK14 | *E. coli* W3110, G1P4 negative secretion, encoding pSKP05/pACYC184 | This study |
| SK15 | *E. coli* W3110, G1P5 positive secretion, encoding pSKP06/pSK01 | This study |

| | | |
|---|---|---|
| SK16 | *E. coli* W3110, G1P5 negative secretion, encoding pSKP06/pACYC184 | This study |
| SK17 | *E. coli* W3110, G1P6 positive secretion, encoding pSKP07/pSK01 | This study |
| SK18 | *E. coli* W3110, G1P6 negative secretion, encoding pSKP07/pACYC184 | This study |
| SK19 | *E. coli* W3110, G1P7 positive secretion, encoding pSKP08/pSK01 | This study |
| SK20 | *E. coli* W3110, G1P7 negative secretion, encoding pSKP08/pACYC184 | This study |
| SK21 | *E. coli* W3110, G1P8 positive secretion, encoding pSKP09/pSK01 | This study |
| SK22 | *E. coli* W3110, G1P8 negative secretion, encoding pSKP09/pACYC184 | This study |
| SK23 | *E. coli* W3110, G1P9 positive secretion, encoding pSKP10/pSK01 | This study |
| SK24 | *E. coli* W3110, G1P9 negative secretion, encoding pSKP10/pACYC184 | This study |
| SK25 | *E. coli* W3110, G1P10 positive secretion, encoding pSKP11/pSK01 | This study |
| SK26 | *E. coli* W3110, G1P10 negative secretion, encoding pSKP11/pACYC184 | This study |
| SK27 | *E. coli* W3110, G2P1 positive secretion, encoding pSKP12/pSK01 | This study |
| SK28 | *E. coli* W3110, G2P1 negative secretion, encoding pSKP12/pACYC184 | This study |
| SK29 | *E. coli* W3110, G2P2 positive secretion, encoding pSKP13/pSK01 | This study |
| SK30 | *E. coli* W3110, G2P2 negative secretion, encoding pSKP13/pACYC184 | This study |
| SK31 | *E. coli* W3110, G2P3 positive secretion, encoding pSKP14/pSK01 | This study |
| SK32 | *E. coli* W3110, G2P3 negative secretion, encoding pSKP14/pACYC184 | This study |
| SK33 | *E. coli* W3110, G2P4 positive secretion, encoding pSKP15/pSK01 | This study |
| SK34 | *E. coli* W3110, G2P4 negative secretion, encoding pSKP15/pACYC184 | This study |
| SK35 | *E. coli* W3110, G2P5 positive secretion, encoding pSKP16/pSK01 | This study |
| SK36 | *E. coli* W3110, G2P5 negative secretion, encoding pSKP16/pACYC184 | This study |
| SK37 | *E. coli* W3110, G2P6 positive secretion, encoding pSKP17/pSK01 | This study |
| SK38 | *E. coli* W3110, G2P6 negative secretion, encoding pSKP17/pACYC184 | This study |
| SK39 | *E. coli* W3110, G2P7 positive secretion, encoding pSKP18/pSK01 | This study |
| SK40 | *E. coli* W3110, G2P7 negative secretion, encoding pSKP18/pACYC184 | This study |
| SK41 | *E. coli* W3110, G2P8 positive secretion, encoding pSKP19/pSK01 | This study |
| SK42 | *E. coli* W3110, G2P8 negative secretion, encoding pSKP19/pACYC184 | This study |

| | | |
|---|---|---|
| SK43 | *E. coli* W3110, G2P9 positive secretion, encoding pSKP20/pSK01 | This study |
| SK44 | *E. coli* W3110, G2P9 negative secretion, encoding pSKP20/pACYC184 | This study |
| SK45 | *E. coli* W3110, G2P10 positive secretion, encoding pSKP21/pSK01 | This study |
| SK46 | *E. coli* W3110, G2P10 negative secretion, encoding pSKP21/pACYC184 | This study |
| SK47 | *E. coli* W3110, G3P1 positive secretion, encoding pSKP22/pSK01 | This study |
| SK48 | *E. coli* W3110, G3P1 negative secretion, encoding pSKP22/pACYC184 | This study |
| SK49 | *E. coli* W3110, G3P2 positive secretion, encoding pSKP23/pSK01 | This study |
| SK50 | *E. coli* W3110, G3P2 negative secretion, encoding pSKP23/pACYC184 | This study |
| SK51 | *E. coli* W3110, G3P3 positive secretion, encoding pSKP24/pSK01 | This study |
| SK52 | *E. coli* W3110, G3P3 negative secretion, encoding pSKP24/pACYC184 | This study |
| SK53 | *E. coli* W3110, G3P4 positive secretion, encoding pSKP25/pSK01 | This study |
| SK54 | *E. coli* W3110, G3P4 negative secretion, encoding pSKP25/pACYC184 | This study |
| SK55 | *E. coli* W3110, G3P5 positive secretion, encoding pSKP26/pSK01 | This study |
| SK56 | *E. coli* W3110, G3P5 negative secretion, encoding pSKP26/pACYC184 | This study |
| SK57 | *E. coli* W3110, G3P6 positive secretion, encoding pSKP27/pSK01 | This study |
| SK58 | *E. coli* W3110, G3P6 negative secretion, encoding pSKP27/pACYC184 | This study |
| SK59 | *E. coli* W3110, G3P7 positive secretion, encoding pSKP28/pSK01 | This study |
| SK60 | *E. coli* W3110, G3P7 negative secretion, encoding pSKP28/pACYC184 | This study |
| SK61 | *E. coli* W3110, G3P8 positive secretion, encoding pSKP29/pSK01 | This study |
| SK62 | *E. coli* W3110, G3P8 negative secretion, encoding pSKP29/pACYC184 | This study |
| SK63 | *E. coli* W3110, G3P9 positive secretion, encoding pSKP30/pSK01 | This study |
| SK64 | *E. coli* W3110, G3P9 negative secretion, encoding pSKP30/pACYC184 | This study |
| SK65 | *E. coli* W3110, G3P10 positive secretion, encoding pSKP31/pSK01 | This study |
| SK66 | *E. coli* W3110, G3P10 negative secretion, encoding pSKP31/pACYC184 | This study |
| SK67 | *E. coli* W3110, G4P1 positive secretion, encoding pSKP32/pSK01 | This study |
| SK68 | *E. coli* W3110, G4P1 negative secretion, encoding pSKP32/pACYC184 | This study |
| SK69 | *E. coli* W3110, G4P2 positive secretion, encoding pSKP33/pSK01 | This study |

| | | |
|---|---|---|
| SK70 | *E. coli* W3110, G4P2 negative secretion, encoding pSKP33/pACYC184 | This study |
| SK71 | *E. coli* W3110, G4P3 positive secretion, encoding pSKP34/pSK01 | This study |
| SK72 | *E. coli* W3110, G4P3 negative secretion, encoding pSKP34/pACYC184 | This study |
| SK73 | *E. coli* W3110, G4P4 positive secretion, encoding pSKP35/pSK01 | This study |
| SK74 | *E. coli* W3110, G4P4 negative secretion, encoding pSKP35/pACYC184 | This study |
| SK75 | *E. coli* W3110, G4P5 positive secretion, encoding pSKP36/pSK01 | This study |
| SK76 | *E. coli* W3110, G4P5 negative secretion, encoding pSKP36/pACYC184 | This study |
| SK77 | *E. coli* W3110, G4P6 positive secretion, encoding pSKP37/pSK01 | This study |
| SK78 | *E. coli* W3110, G4P6 negative secretion, encoding pSKP37/pACYC184 | This study |
| SK79 | *E. coli* W3110, G4P7 positive secretion, encoding pSKP38/pSK01 | This study |
| SK80 | *E. coli* W3110, G4P7 negative secretion, encoding pSKP38/pACYC184 | This study |
| SK81 | *E. coli* W3110, G4P8 positive secretion, encoding pSKP39/pSK01 | This study |
| SK82 | *E. coli* W3110, G4P8 negative secretion, encoding pSKP39/pACYC184 | This study |
| SK83 | *E. coli* W3110, G4P9 positive secretion, encoding pSKP40/pSK01 | This study |
| SK84 | *E. coli* W3110, G4P9 negative secretion, encoding pSKP40/pACYC184 | This study |
| SK85 | *E. coli* W3110, G4P10 positive secretion, encoding pSKP41/pSK01 | This study |
| SK86 | *E. coli* W3110, G4P10 negative secretion, encoding pSKP41/pACYC184 | This study |
| SK87 | *E. coli* DH5α, codon optimized G1P6 positive secretion, encoding pSKP42/pSK01 | This study |
| SK88 | *E. coli* DH5α, codon optimized G1P6 negative secretion, encoding pSKP42/pACYC184 | This study |
| SK89 | *E. coli* DH5α, G1P6_2X positive secretion, encoding pSKP43/pSK01 | This study |
| SK90 | *E. coli* DH5α, G1P6 _2X negative secretion, encoding pSKP43/pACYC184 | This study |
| SK91 | *E. coli* DH5α, G3P2 positive secretion, encoding pSKP23/pSK01 | This study |
| SK92 | *E. coli* DH5α, G3P2 negative secretion, encoding pSKP23/pACYC184 | This study |
| SK93 | *E. coli* DH5α, G3P2_2X positive secretion, encoding pSKP44/pSK01 | This study |
| SK94 | *E. coli* DH5α, G3P2_2X negative secretion, encoding pSKP44/pACYC184 | This study |
| SK95 | *E. coli* W3110, Pediocin PA-1 positive secretion, encoding pSKP45/pSK01 | This study |

| | | |
|---|---|---|
| SK96 | *E. coli* W3110, Pediocin PA-1 negative secretion, encoding pSKP45/pACYC184 | This study |
| SK97 | *E. coli* W3110, Pediocin PA-1 protease-deficient secretion, encoding pSKP45/pSK02 | This study |
| SK98 | *E. coli* W3110, α-factor positive secretion, encoding pSKP46/pSK01 | This study |
| SK99 | *E. coli* W3110, α-factor negative secretion, encoding pSKP46/pACYC184 | This study |
| SK100 | *E. coli* W3110, α-factor protease-deficient secretion, encoding pSKP46/pSK02 | This study |
| SK101 | *E. coli* W3110, eglin C positive secretion, encoding pSKP47/pSK01 | This study |
| SK102 | *E. coli* W3110, eglin C protease-deficient secretion, encoding pSKP47/pSK02 | This study |
| SK103 | *E. coli* W3110, EGF positive secretion, encoding pSKP48/pSK01 | This study |
| SK104 | *E. coli* W3110, EGF negative secretion, encoding pSKP48/pACYC184 | This study |
| SK105 | *E. coli* Nissle 1917, Pediocin PA-1 positive secretion, encoding pSKP49 | This study |
| SK106 | *E. coli* Nissle 1917, Pediocin PA-1 negative secretion, encoding pSKP50 | This study |
| SK107 | *Salmonella enterica* Ty21a, Pediocin PA-1 positive secretion, encoding pSKP49 | This study |
| SK108 | *Salmonella enterica* Ty21a, Pediocin PA-1 negative secretion, encoding pSKP50 | This study |
| SK109 | *Vibrio cholerae* CVD103-HgR, Pediocin PA-1 positive secretion, encoding pSKP49 | This study |
| SK110 | *Vibrio cholerae* CVD103-HgR, Pediocin PA-1 negative secretion, encoding pSKP50 | This study |
| SK111 | *E. coli* BL21(DE3), empty vector, encoding pBAD18-Km/pACYC184 | This study |
| SK112 | *E. coli* BL21(DE3), G1P5 positive secretion, encoding pSKP06/pSK01 | This study |
| SK113 | *E. coli* BL21(DE3), G1P10 positive secretion, encoding pSKP11/pSK01 | This study |
| SK114 | *E. coli* BL21(DE3), G2P6 positive secretion, encoding pSKP17/pSK01 | This study |
| SK115 | *E. coli* BL21(DE3), G2P7 positive secretion, encoding pSKP18/pSK01 | This study |
| SK116 | *E. coli* BL21(DE3), G3P3 positive secretion, encoding pSKP24/pSK01 | This study |
| SK117 | *E. coli* BL21(DE3), G3P5 positive secretion, encoding pSKP26/pSK01 | This study |
| SK118 | *E. coli* BL21(DE3), G1P5 negative secretion, encoding pSKP06/pACYC184 | This study |
| SK119 | *E. coli* BL21(DE3), G1P10 negative secretion, encoding pSKP11/pACYC184 | This study |
| SK120 | *E. coli* BL21(DE3), G2P6 negative secretion, encoding pSKP17/pACYC184 | This study |
| SK121 | *E. coli* BL21(DE3), G2P7 negative secretion, encoding pSKP18/pACYC184 | This study |

| | | |
|---|---|---|
| SK122 | *E. coli* BL21(DE3), G3P3 negative secretion, encoding pSKP24/pACYC184 | This study |
| SK123 | *E. coli* BL21(DE3), G3P5 negative secretion, encoding pSKP26/pACYC184 | This study |
| SK124 | *E. coli* BL21(DE3), EGF_strep positive secretion, encoding pSKP51/pSK01 | This study |
| SK125 | *E. coli* BL21(DE3), Eglin C_strep positive secretion, encoding pSKP52/pSK01 | This study |

**Table S4. Primers and gBlocks.**

| Type | Name | Sequence | Usage |
|---|---|---|---|
| primer | random_20mer_V5 | ATAGAGCTCGAATTCAGGAGGAAACGATGAGAACTCTGACTCTAAATGAATTAGATTCTGTTTCTGGTGGTNNKNNKNNKNNKNNKNNKNNKNNKNNKNNKNNKNNKNNKNNKNNKNNKNNKNNKNNKNNKGGAGGAGGTAAACCTATTCCTAATCCTCTCCTAGGTTTAGATTCTACTTAAGTCGACAGGAGGAAACGA | template for generating group2 peptide ORFs |
| primer | cvaC15_induce_F | ATAGAGCTCGAATTCAGGAGGAAACGATGAGAACTCTGACTCTAAATG | amplifying DNA containing MccV signal peptide sequences |
| primer | V5_R | TATGTCGACTTAAGTAGAATCTAAACCTAGGAGAGG | amplifying DNA containing V5-tag sequences |
| primer | cvaB_mutation_F | CATCAGACGGAGACCGCTGAATCTGGACTG | to make pSK02 |
| primer | cvaB_mutation_R | GAGCCCGGGTGGCCATCC | to make pSK02 |
| primer | pBAD_vecF | GTCTGCTTACATAAACAGTAATACAAGGGGTG | to delete Sfo I site of pBAD_cvi_cvaC |
| primer | pBAD_vecR | GCGCCACAGGTGCGGTTGCTATCTCCTTGCTGCCTCGCG | to delete Sfo I site of pBAD_cvi_cvaC |
| primer | pBAD_fragF | GCGCGAGGCAGCAAGGAGATAGCAACCGCACCTGTGG | to delete Sfo I site of pBAD_cvi_cvaC |
| primer | pBAD_fragR | GCTCATAACACCCCTTGTATTACTGTTTATGTAAGCAGACAGTTTTATTGTTCATGA | to delete Sfo I site of pBAD_cvi_cvaC |
| gBlock | pBAD_MCS_2 | ATAGAGCTCGAATTCAGGAGGAAACGATGAGAACTCTGACTCTAAATGAATTAGATTCTGTTTCTGGCGCCGGTACCCGGGGATCCTCTAGAGTCGACCTGCAGGCATGCAAGCTTGGCTGTTTTGGCGGATGAGAGAAGATTTTCAGCCTGATACAGATTAAATCAGAACGCAGAAGCGGTCTGATAAAACAGAATTTGCCTGGCGGCAGTAGCGCGGTGGTCCCACCTGACCCCATGCCGAACTCAGAAGTGAAACGCCGTAGCGCCGATGGTAGTGTGGACTAGTGGTCTCCCCATGCATA | template for generating pSK00 (MccV signal peptide, sfoI site containing) |
| primer | pBAD_MCS_2_R | TATGCATGGGGAGACCACTAGTCCACACT | to generate pSK00 or amplify the region |
| gBlock | EglinC | GAGCTCGAATTCAGGAGGAAACGATGAGAACTCTGACTCTAAATGAATTAGATTCTGTTTCTGGTGGTACCGAATTTGGCAGCGAACTGAAAAGCTTTCCGGAAGTGGTGGGCAAAACCGTGGATCAGGCGCGCGAATATTTTACCCTGCATTATCCGCAGTATGATGTGTATTTTCTGCCGGAAGGCAGCCCGGTGACCCTGGATCTGCGCTATAACCGCGTGCGCGTGTTTTATAACCCGGG | to clone eglinC into pBAD |

| | | | |
|---|---|---|---|
| | | CACCAACGTGGTGAACCATGTGCCGCATGTGGGCGGAGGAGGTAAACCTATTCCTAATCCTCTCCTAGGTTTAGATTCTACTTAAAAGTCGAC | |
| primer | EglinC_R | ATAGTCGACTTAGCCCACATGCGGCACATGGT | to amplify eglinC |
| primer | pBAD_N10mer | ATCCCGGGNNKNNKNNKNNKNNKNNKNNKNNKNNKNNKGGAGGAGGTAAACCTATTCCTAATCCTCTCCTAGGT | to generating group1 peptide ORFs |
| primer | G3P2_bbs1 | ATAGAGCTCGAAGACCATGGCGTGCAGGGC | to generate BbsI site containing G3P2 |
| primer | G3P2_bbs2 | ATGAAGACAAGCCATGGTTCTGTTCATAGCTATCATCGCGG | to generate G3P2_2X via SacI and BbsI cloning |
| gBlock | Pediocin PA-1 | AAATACTACGGCAATGGGGTGACCTGTGGGAAACATTCCTGCTCCGTTGACTGGGGGAAAGCGACCACCTGTATCATCAATAACGGAGCGATGGCCTGGGCTACGGGCGGTCACCAGGGCAATCACAAGTGT | template for Pediocin PA-1 |
| primer | Pediocin_F | ATAGAGCTCAGGAGGAAACGATGAGAACTCTGACTCTAAATGAATTAGATTCTGTTTCTGGTGGTAAATACTACGGCAATGGTGTAACG | amplifying Pediocin PA-1 |
| primer | Pediocin_F | ATATCTAGATTAGCACTTATGATTTCCCTGGTGGC | amplifying Pediocin PA-1 |
| primer | alpha_F | ATCCCGGGTGGCACTGGCTGCAGCTGAAACCGGGTCAGCCGATGTACTAAGGTACCTGGGGATCCTCTAGAG | amplifying alpha-factor |
| gBlock | EGF | ATACCCGGGAACAGCGATAGCGAATGCCCGCTGAGCCATGATGGCTATTGCCTGCATGATGGCGTGTGCATGTATATTGAAGCGCTGGATAAATATGCGTGCAACTGCGTGGTGGGCTATATTGGCGAACGCTGCCAGTATCGCGATCTGAAATGGTGGGAACTGCGCTAAGTCGACATA | template for EGF |
| primer | EGF_F | ATACCCGGGAACAGCGATAG | to generate EGF |
| primer | EGF_R | TATGTCGACTTAGCGCAGTTCCC | to generate EGF |
| gBlock | Random1 | ACACTTTGCTATGCCCGGGTATATGTGCTGGCATCAGGCGATTAACGTGATGGAACCGTGCGCGCAGTTTTATCAGGATATTGTGCTGAGCAGCCGCGTGCAGTGGCAGGATATGGATATGAGCATGCCGCGCCTGTATAAAATGCAGTATGTGGCGAAAAGCCATTTTAGCATTATGTATTTTATTCATCGCGAAGATATTCAGAGCAGCGGCTGCTGCGATTGCAACTGCAACCCGGTGCGCAAAATTTTTTGCACCCGCTATGTGGAAATGGTGGATTGCGGCTATTGGCGCCATTTTTGGCTG | template for G3P1 and G4P1 |

| | | | |
|---|---|---|---|
| | | CCGCCGGAAGAAGGAGGAGGTAAACCTATT CCTAATCCCGGGCTGAGCAACGGCCAGTGC AACATGCATTGCTGCCCGTGCCTGGAATATC AGGATTATCATAAACATTATAGCAACACCGA AAGCTTTAAACAGCTGGTGTGGATGACCCAT ATTTGCGATAACTATGCGCTGAGCCATCGCG CGAAATGGGGAGGACGTGATGTCTTC | |
| gBlock | Random2 | ACACTTTGCTATGCCCGGGTTTATTATTTATA TTCCGAAAGTGTATGATTGGTATGCGGCGAG CATGGGCTGCACCTATCCGGATGGCCTGGG CTTTCGCATGGTGAGCATGCGCGTGATTTG GTATGCGTGGTATGTGTATAGCTGCTATACC GCGGTGGAAACCAACAAATTTGGCGATTGC GGCTGCGGCAACTATAACCATCCGGATCAG GAACAGATTCGCTATAAAACCAGCGAACAGG AAAGCACCAAAATGTTTTTTCGCTGGGCGCC GGAATGGCAGAACAACCACAGGCACCACCT GATGGTGGAAATGGGAGGAGGTAAACCTAT TCCTAATCCCGGGCATGGCGTGCAGGGCAT TAACATTGAAAAACAGCCGAAACGCAACAAC CCGGAAAACGAACAGACCCGCATGAAAATG CGCCAGGAACGCGATTGGAGCTGCTTTGCG ATGTTTCATGCGATTACCCGCGATGATAGCT ATGAACAGAAC | template for G3P2 and G4P2 |
| gBlock | Random3 | ACACTTTGCTATGCCCGGGCACAGTTATTGC GAATGGCTGATGGCGAAAATTCTGAGCATG GGCGAACAGTGGTGGCATAAATATTATTTTG GCCTGCGCCATCAGTTTAACGTGAGCAAAG GCTATCATTTTAACACCAGCTTTGATTTTCAT CGCTGCGGCGAACCGAAAAACAACTATTATG CGCGCACCCATTGCGAAAAAGAATATAGCAA CGATGATGTGCATCCGCGCCAGAACAGCAT GGGCCGCTGGGGCGCGGAATGGCCGGCGC TGATTTTTAAACCGCTGTTTGGCATTAACAAA TGGGGCAACTTTGGAGGAGGTAAACCTATTC CTAATCCCGGGGAAACCTGCCATTGGATGG AACTGCATATTCCGCTGTTTGAAACCGATAG CTTTAAACCGTATGATCCGAAAAGCCTGGAT AGCGGCCATTGCCTGTATTATGGCTTTTTTTT TAAATATATTGGCGGCCTGCATGCGATGTGC ATGTATGGAGGACGTGATGTCTTC | template for G3P3 and G4P3 |
| gBlock | Random4 | ACACTTTGCTATGCCCGGGAGCGGCATTTAT TGCCAGATTACCCGCTGGGTGCATCCGTTTA GCGAAAGCACCCAGAACATGGATAACATGG CGAACACCAAACGCAAACCGCAGTGGCATT ATCCGCGCCGCCATCAGCATAAAGAAGCGC | template for G3P4 and G4P4 |

| | | | |
|---|---|---|---|
| | | AGTTTGTGTTTTTTGGCATTGCGCGCGGCTG GCCGTTTCAGTTTTTTGAACAGGGCTGGGTG ACCATTGATGTGCATGAAGAACATCTGTGGA TTTTTTGCTTTGGCGAACGCAACATGGAACA TGGCAACGTGGAACGCGCGCCGACCATTAG CAGCCGCATTAAAGGAGGAGGTAAACCTATT CCTAATCCCGGGGTGCAGATGATTTGGGGC TTTTGCAGCGGCTGGCCGATGATTACCTGGT ATGCGATGATGTTTGCGCATATTCAGTGGGC GTTTTGGAACACCAAAATTAGCAGCCGCGG CTGGGAATTTATGGCGAGCGCGTGGCCGGA ATATTTTGTGGGAGGACGTGATGTCTTC | |
| gBlock | Random5 | ACACTTTGCTATGCCCGGGAAACATTGGAAA CAGCGCCATGTGTATTATCGCCAGGCGGTG TATCAGCAGCAGAAATGCCAGATGTATAACG TGCCGTATAGCCCGGCGAGCGGCTGCTTTA ACTGCCAGCCGAACTGCCATCGCAAAGATTT TGATTGGCGCGATCATACCGGCTATTGCTTT AAAATGCTGCAGTTTCATAACTTTATTACCCC GAAACGCTGCACCGAAAGCGCGCTGTTTGC GGGCCAGGAATGCCGCCCGGATTTTGCGCA GGAAAGCGGCGCGCAGAAAGATTTTAGCGG CATGCATCCGGGCGGAGGAGGTAAACCTAT TCCTAATCCCGGGATTATGGAATGCAGCACC CTGATTTGCACCACCCTGGATTGCTTTATTT GCGTGCAGGGCCAGTGGCGCTATTGCATGT GGAACCAGTGCTGGTGCGTGTGCATGAACT GGACCAAAACCAACTGCGCGCAGTATAGCG TGGCGAAACATGGAGGACGTGATGTCTTC | template for G3P5 and G4P5 |
| gBlock | Random6 | ACACTTTGCTATGCCCGGGTTTAACCCGGAA AGCAGCCATCAGCCGAGCACCATTCCGAAA AGCCATTTTCGCATTATTTGCCATTTTGTGCG CGATTGGCATTTTCCGTGCGGCAGCTGGAC CTTTAGCGTGGTGGATATTTATTTTTGCGAAA TGTATACCACCCTGGGCAACCCGCATGGCT TTGTGATTTGCTGCACCTATGGCAGCCAGTA TAGCGGCGATAACCGCTGCGCGGATAAACT GGAACGCCATCCGGCGATGATGGAAAACAC CTATGGCTGGCATGGCCATACCAGCGCGGG CCTGGCGCAGCCGGGAGGAGGTAAACCTAT TCCTAATCCCGGGATTATTTGCTGGTGGCCG CATCCGCAGTGCTGCTGGAACTTTGAATATT GCTTTCGCAAAAACTATCTGACCTGCTTTCA GTGCAGCGAACAGTATAGCACCACCTTTGC GCCGGTGAACGCGTTTCCGACCATTTGGCA GATTATTTATGGAGGACGTGATGTCTTC | template for G3P6 and G4P6 |

| | | | |
|---|---|---|---|
| gBlock | Random7 | ACACTTTGCTATGCCCGGGGATAAAATGGAACCGCAGTGGAACCATAGCCCGCGCTGCTTTGCGAGCCTGTGCTGCGGCGGCAGCCATACCATGATGGCGTGGAACCATGTGATTAGCTGGAAAGGCCGCGATCTGGTGATTGGCGTGAACCGCCATTGCGCGACCCCGCCGCATAGCCAGTTTTGGCATAACGCGTGGTGGCAGGGCTTTATTAAACATGTGATTGAAACCCCGCGCCTGGGCAACATGGCGAAAAACATGCAGTGCTTTGCGGCGTGCGCGCTGGCGGTGGCGTTTCCGGTGGATATTAGCCAGCTGGGAGGAGGTAAACCTATTCCTAATCCCGGGGGCAGCGTGATTGTGCTGGATTGGGTGCAGGGCACCAACCTGGTGCATAAACAGCATAACACCATTAACCGCCGCCATCATCATAGCCATCAGGAAATGTATCCGTGGCCGACCGTGTTTGAAATGCATGGGAACCGCATTATTGAAGAAGGAGGACGTGATGTCTTC | template for G3P7 and G4P7 |
| gBlock | Random8 | ACACTTTGCTATGCCCGGGCGCGGCAAATGGATGATGTGCAACACCCATATGGCGCATTGGAACACCTATTTTTGGGATGGCAGCGCGGTGCATCTGACCGATGATTTTTATCGCAACGGCCCGGCGAAAAGCTATAACCTGTTTATGGTGCAGAACCATAAAGAAAGCCGCCATAACTATAAAGTGTGCTTTTTTTGCTATCTGATTACCACCTATATTACCATTGCGAAACATAAACGCATGAACGAAAACTATTGGTGGATGGCGCAGGTGTATCTGAAATTTGTGCGCTGGCATGCGCGCAACTGCTGCTATGCGGGAGGAGGTAAACCTATTCCTAATCCCGGGCATGTGCCGAAATGGTGGTATAAAGGCTTTGATTGGACCACCCAGGTGTGGCCGTATGCGGCGATGCTGGGCTTTATTAACGCGCATCATCTGGATACCGTGATGATTAAAATTCATCTGGGCGCGTTTCATAACTGCGATTGGGTGGAAGGAGGACGTGATGTCTTC | template for G3P8 and G4P8 |
| gBlock | Random9 | ACACTTTGCTATGCCCGGGGTGATGCAGGGCGATACCTGGTGGGAATGCGAACCGAGCGAAGCGGAAATTCAGATGCTGTATTGGCCGTGGGGCAGCCAGAAAGATCCGATTGATTGGGCGTATCTGTGCGATACCTGGAAATATACCGGCGATCTGTGCAGCGGCGGCCCGGAACAGCCGGATGAACATCGCATTCATGATGCGATTGGCCGCGCGTTTTATCGCCCGTGCCCGAGCCTGAACATGTATTATCTGAGCCAGCGCTGGGCGATTTTTGATACCCATAACAGCCTGGCGGCGGGCAGCTATTGCTTTATGGGAGGAGGTAAACC | template for G3P9 and G4P9 |

| | | | |
|---|---|---|---|
| | | TATTCCTAATCCCGGGGTGACCACCTTTAAA<br>CTGTGGGCGAAAGCGCTGGTGGCGTTTATG<br>TATGATGCGGATCATCATGTGAACGATTTTC<br>TGCCGACCCTGTATCGCGTGTATACCACCAT<br>GAACATTTGGCATTTTAAACATAAATGCCCG<br>TGCACCAGCTATGGAGGACGTGATGTCTTC | |
| gBlock | Random10 | ACACTTTGCTATGCCCGGGACCGGCCGCCA<br>GGTGACCGAAGTGACCGTGTGGCATGCGCT<br>GACCACCATTTGCGGCATTAGCGAACTGGA<br>ATTTACCTATGGCGCGTGCCCGATGTGGGA<br>AAACATGGAACTGGAAAAATTTAGCGGCAAC<br>GTGTGCTATGAACTGCAGGATCATTGCTTTT<br>GCGATTGGTGGCAGTATACCGAACGCTGCC<br>TGGAAAACCTGCCGATGATTGAACTGCCGAT<br>TCAGTGGAAACCGTTTACCCTGCATGAATGG<br>TGGATTATTGGCCGCTGCCCGCTGACCATTA<br>TGAACAGCTGGGCGGGAGGAGGTAAACCTA<br>TTCCTAATCCCGGGGATGATGATGATGAACT<br>GCTGCGCCTGTGCGATAACATTACCTTTTTT<br>ATGATGTGCATTCATGAATTTACCATGAAAC<br>CGTGGTTTAAAACCATTTGGTTTCTGATGTG<br>CTGGAACGCGTTTCTGAACCAGGGCAGCAA<br>TAGCACCCATGGAGGACGTGATGTCTTC | template for G3P10 and G4P10 |
| primer | 100mer_F | ACACTTTGCTATGCCCGGG | amplifying group 4 random peptides |
| primer | 100mer_R | GGGATTAGGAATAGGTTTACCTCCTCC | amplifying group 4 random peptides |
| primer | 50mer_F | GAGGTAAACCTATTCCTAATCCCGGG | amplifying group 3 random peptides |
| primer | 50mer_R1 | AGGAATAGGTTTACCTCCTCCCCATTTCGCG<br>CGATGGC | amplifying G3P1 |
| primer | 50mer_R2 | AGGAATAGGTTTACCTCCTCCGTTCTGTTCA<br>TAGCTATCATCGCGG | amplifying G3P2 |
| primer | 50mer_R3 | AGGAATAGGTTTACCTCCTCCATACATGCAC<br>ATCGCATGCAG | amplifying G3P3 |
| primer | 50mer_R4 | AGGAATAGGTTTACCTCCTCCCACAAAATAT<br>TCCGGCCACGC | amplifying G3P4 |
| primer | 50mer_R5 | AGGAATAGGTTTACCTCCTCCATGTTTCGCC<br>ACGCTATACTGC | amplifying G3P5 |
| primer | 50mer_R6 | AGGAATAGGTTTACCTCCTCCATAAATAATCT<br>GCCAAATGGTCGGAAACG | amplifying G3P6 |
| primer | 50mer_R7 | AGGAATAGGTTTACCTCCTCCTTCTTCAATAA<br>TGCGGTTCCCATGC | amplifying G3P7 |
| primer | 50mer_R8 | AGGAATAGGTTTACCTCCTCCTTCCACCCAA<br>TCGCAGTTATGAAA | amplifying G3P8 |

| | | | |
|---|---|---|---|
| primer | 50mer_R9 | AGGAATAGGTTTACCTCCTCCATAGCTGGTGCACGGGCAT | amplifying G3P9 |
| primer | 50mer_R10 | AGGAATAGGTTTACCTCCTCCATGGGTGCTATTGCTGCCCT | amplifying G3P10 |
| primer | G1P6_opti_F | ATCCCGGGCTGTCTATGTCTATCTGCATGCGTCCGAAAGGAGGAGGTAAACCTATTCCTA | amplifying G1P6 with *E.coli* optimized codon |
| primer | G1P6_2X_opti_F | ATCCCGGGCTGTCTATGTCTATCTGCATGCGTCCGAAACTGTCTATGTCTATCTGCATGCGTCCGAAAGGAGGAGGTAAACCTATTCCTA | amplifying G1P6_2X with *E.coli* optimized codon |
| primer | cvi_cvaC_pBAD_F | GGTACCAGGAGGAAACGATGGATAGAAAAAGAACAAAATTAGAGTTGTTATTTGC | first cloning of *cvi* and *cvaC* into pBAD180-Km |
| primer | pBAD_cvi_F | GGTGGTGAATTCAGGAGGAAACGATGGATAGAAA | amplifying *cvi* |
| primer | pBAD_cvi_R | GCGTGGTACCTCATTTAGAGTCAGAGTTC | amplifying *cvi* |
| primer | pBAD_cvaC_F | GGTGGTGGTACCATGAGAACTCTGACTCTAAAT | amplifying *cvaC* |
| primer | pBAD_cvaC_R | GGCGGCGTCGACTCTAGATTATAAACAAACATCACT | amplifying *cvaC* |
| primer | MccV_V5_R | CAGGTCGACTTAAGTAGAATCTAAACCTAGGAGAGGATTAGGAATAGGTTTACCTCCTCCTAAACAAACATCACTAAGATTATTTGGACT | to generate MccV_V5 |
| primer | CvaAB_F | ATATCTAGATTTCAGTGCAATTTATCTCTTCAAATGTAGCACCTGAAGTCAGCCCCATACGATATAAGTTGTAATTCTCATGTTTGACAGCTTATCATCGATAAGCTTTAATGCGGTAGTTTATCACAGTTAAATTGCTAACGCAGTCAGGCACCGTGTAGGAGGAAACGATGTTTCGCCAGGATGCTTTAGAAAAC | amplifying *cvaA*/*cvaB* with pTc promoter |
| primer | CvaAB_R | ATACCTGAGGGTATTATTTAATATAAGAAAGAACAGTTATTGGACAATCCAC | amplifying *cvaA*/*cvaB* |
| primer | CvaAB_F2 | ATAGAGCTCTGGGTACCCGGGGATCCTCTAGAGTCGACAGGAGGAAACGATG | amplifying *cvaA*/*cvaB* to construct pSK03 |
| primer | CvaAB_R2 | ATAGCATGCCTGCAGTTAAATAGAAATAACTC | amplifying *cvaA*/*cvaB* to construct pSK03 |
| primer | EGF_strep_R | ATAGTCGACTTATTTTTCGAACTGCGGGTGAGACCATCCTCCGCGCAGTTCCCACCATTTCAGATC | to generate EGF_strep |
| primer | Eglin C_strep_R | ATAGTCGACTTATTTTTCGAACTGCGGGTGAGACCATCCTCCGCCCACATGCGGCACATGGTT | to generate EglinC_strep |

**cvaC*: encodes microcin V (MccV)

**Table S5. Plasmids**

| Name | Description | Usage |
|---|---|---|
| pACYC184 | Cm$^R$, Tet$^R$ | Backbone plasmid |
| pBAD18-Km | Kan$^R$ | Backbone plasmid |
| pMMB67EH | Amp$^R$ | Backbone plasmid |
| pBR322 | Amp$^R$ | Template |
| pHK22 | Amp$^R$ | Template |
| pSK00 | pBAD18-Km derived plasmid, Kan$^R$ | POI expression |
| pSK01 | pACYC184 derived plasmid, Cm$^R$ | CvaA/CvaB expression |
| pSK02 | pACYC184 derived plasmid, Cm$^R$ | CvaA/CvaB C32S expression |
| pSK03 | pMMB67EH derived plasmid, Amp$^R$ | POI, CvaA/CvaB expression |
| pSKP00 | pBAD18-Km derived plasmid, Kan$^R$ | Cvi, MccV expression |
| pSKP01 | pBAD18-Km derived plasmid, Kan$^R$ | Cvi, MccV_V5 expression |
| pSKP02 | pBAD18-Km derived plasmid, Kan$^R$ | G1P1 expression |
| pSKP03 | pBAD18-Km derived plasmid, Kan$^R$ | G1P2 expression |
| pSKP04 | pBAD18-Km derived plasmid, Kan$^R$ | G1P3 expression |
| pSKP05 | pBAD18-Km derived plasmid, Kan$^R$ | G1P4 expression |
| pSKP06 | pMMB67EH derived plasmid, Amp$^R$ | G1P5 expression |
| pSKP07 | pBAD18-Km derived plasmid, Kan$^R$ | G1P6 expression |
| pSKP08 | pBAD18-Km derived plasmid, Kan$^R$ | G1P7 expression |
| pSKP09 | pBAD18-Km derived plasmid, Kan$^R$ | G1P8 expression |
| pSKP10 | pBAD18-Km derived plasmid, Kan$^R$ | G1P9 expression |
| pSKP11 | pBAD18-Km derived plasmid, Kan$^R$ | G1P10 expression |
| pSKP12 | pBAD18-Km derived plasmid, Kan$^R$ | G2P1 expression |
| pSKP13 | pBAD18-Km derived plasmid, Kan$^R$ | G2P2 expression |
| pSKP14 | pBAD18-Km derived plasmid, Kan$^R$ | G2P3 expression |
| pSKP15 | pBAD18-Km derived plasmid, Kan$^R$ | G2P4 expression |
| pSKP16 | pBAD18-Km derived plasmid, Kan$^R$ | G2P5 expression |
| pSKP17 | pBAD18-Km derived plasmid, Kan$^R$ | G2P6 expression |
| pSKP18 | pBAD18-Km derived plasmid, Kan$^R$ | G2P7 expression |
| pSKP19 | pBAD18-Km derived plasmid, Kan$^R$ | G2P8 expression |
| pSKP20 | pBAD18-Km derived plasmid, Kan$^R$ | G2P9 expression |
| pSKP21 | pBAD18-Km derived plasmid, Kan$^R$ | G2P10 expression |

| | | |
|---|---|---|
| pSKP22 | pBAD18-Km derived plasmid, $Kan^R$ | G3P1 expression |
| pSKP23 | pBAD18-Km derived plasmid, $Kan^R$ | G3P2 expression |
| pSKP24 | pBAD18-Km derived plasmid, $Kan^R$ | G3P3 expression |
| pSKP25 | pBAD18-Km derived plasmid, $Kan^R$ | G3P4 expression |
| pSKP26 | pBAD18-Km derived plasmid, $Kan^R$ | G3P5 expression |
| pSKP27 | pBAD18-Km derived plasmid, $Kan^R$ | G3P6 expression |
| pSKP28 | pBAD18-Km derived plasmid, $Kan^R$ | G3P7 expression |
| pSKP29 | pBAD18-Km derived plasmid, $Kan^R$ | G3P8 expression |
| pSKP30 | pBAD18-Km derived plasmid, $Kan^R$ | G3P9 expression |
| pSKP31 | pBAD18-Km derived plasmid, $Kan^R$ | G3P10 expression |
| pSKP32 | pBAD18-Km derived plasmid, $Kan^R$ | G4P1 expression |
| pSKP33 | pBAD18-Km derived plasmid, $Kan^R$ | G4P2 expression |
| pSKP34 | pBAD18-Km derived plasmid, $Kan^R$ | G4P3 expression |
| pSKP35 | pBAD18-Km derived plasmid, $Kan^R$ | G4P4 expression |
| pSKP36 | pBAD18-Km derived plasmid, $Kan^R$ | G4P5 expression |
| pSKP37 | pBAD18-Km derived plasmid, $Kan^R$ | G4P6 expression |
| pSKP38 | pBAD18-Km derived plasmid, $Kan^R$ | G4P7 expression |
| pSKP39 | pBAD18-Km derived plasmid, $Kan^R$ | G4P8 expression |
| pSKP40 | pBAD18-Km derived plasmid, $Kan^R$ | G4P9 expression |
| pSKP41 | pBAD18-Km derived plasmid, $Kan^R$ | G4P10 expression |
| pSKP42 | pBAD18-Km derived plasmid, $Kan^R$ | codon-optimized G1P6 expression |
| pSKP43 | pBAD18-Km derived plasmid, $Kan^R$ | G1P6_2X expression |
| pSKP44 | pBAD18-Km derived plasmid, $Kan^R$ | G3P2_2X expression |
| pSKP45 | pBAD18-Km derived plasmid, $Kan^R$ | Pediocin PA-1 expression |
| pSKP46 | pBAD18-Km derived plasmid, $Kan^R$ | α-factor |
| pSKP47 | pBAD18-Km derived plasmid, $Kan^R$ | Eglin C expression |
| pSKP48 | pBAD18-Km derived plasmid, $Kan^R$ | EGF expression |
| pSKP49 | pMMB67EH derived plasmid, $Amp^R$ | Pediocin PA-1, CvaA/CvaB expression |
| pSKP50 | pMMB67EH derived plasmid, $Amp^R$ | Pediocin PA-1 expression |
| pSKP51 | pBAD18-Km derived plasmid, $Kan^R$ | EGF_strep expression |
| pSKP52 | pBAD18-Km derived plasmid, $Kan^R$ | Eglin C_strep expression |
